# Selfish mitochondria exploit nutrient status across different levels of selection

**DOI:** 10.1101/2020.01.30.927202

**Authors:** Bryan L. Gitschlag, Ann T. Tate, Maulik R. Patel

## Abstract

Cooperation and cheating are widespread evolutionary strategies. Competition can simultaneously favor cheating within groups and cooperation between groups. Selfish or cheater mitochondrial DNA (mtDNA) mutants proliferate within hosts while being selected against at the level of host fitness. How does environment govern competition between cooperators and cheaters across different selection levels? Focusing on food availability, we address this question using heteroplasmic *Caenorhabditis elegans*. We show that by promoting germline development, nutrient status provides the niche space for mtDNA variants to compete. However, the within-host advantage of selfish mtDNA requires additional conditions, namely the FoxO transcription factor DAF-16. During food scarcity, DAF-16 mitigates the host fitness cost of the selfish mtDNA. We conclude that food availability, and resilience to food scarcity, govern selfish mtDNA dynamics across the levels of selection. Our study integrates an evolutionary framework with experimentation to identify cellular mechanisms underlying the multilevel selection that characterizes cheater dynamics.

## Introduction

Life is generally organized into a hierarchy of cooperative collectives, with multiple genes collectively composing a genome, different genomes giving rise to a eukaryotic cell, cells giving rise to multicellular organisms, and multicellular organisms forming societies (Wilson and Wilson, 2007). New levels of organization in the hierarchy emerge when natural selection favors a division of labor and the loss of conflict between previously autonomous replicators, giving rise to a cohesive unit upon which selection can further act (Michod et al., 2006; West et al., 2015). Cooperation thus facilitates evolutionary transitions toward larger, more complex systems (Fisher and Regenberg, 2019; Gulli et al., 2019; Michod et al., 2006; West et al., 2015). However, because cooperation requires investing in the fitness of others for mutual benefit, it also creates the conditions that enable the emergence of selfish “cheater” entities. By benefiting from the cooperative contributions of others without reciprocating, cheaters experience a fitness advantage over their cooperative counterparts (Dobata et al., 2009; Ghoul et al., 2014; Strassmann et al., 2000). This advantage can break down at higher levels of biological organization, which rely on cooperation at lower levels (de Vargas Roditi et al., 2013; Ozkaya et al., 2017). Hence, natural selection can favor a trait at one level of the biological hierarchy while favoring a competing trait at a different level, a phenomenon known as multilevel selection (de Vargas Roditi et al., 2013; Wilson and Wilson, 2007). Multilevel selection thus provides an explanation for the paradoxical coexistence of selfishness and cooperation in an evolving population.

Competition over limited resources shapes relative reproductive fitness, and hence Darwinian evolution. By ensuring efficient resource utilization, cooperation can provide an adaptive benefit (Koschwanez et al., 2013; Vanthournout et al., 2016). Accordingly, the availability of resources likely represents an important ecological determinant of the population dynamics between selfish and cooperative entities (Ducasse et al., 2015). Prior studies have indicated that resource scarcity can favor cooperative, group-oriented interactions (Cao et al., 2015; Chisholm and Firtel, 2004; Li and Purugganan, 2011; Pereda et al., 2017), whereas resource abundance can promote selfishness (Chen and Perc, 2014). The link between resource availability and the proliferation of selfish entities can be found at multiple levels of scale, with cancer representing a well-known example. Cancer cells can selfishly exploit nutrient supply, since nutrient abundance stimulates the production of growth factors, increasing cell growth and division (Han et al., 2015; Narita et al., 2019; Tucci, 2012). Conversely, nutrient restriction promotes the conservation of resources through autophagy, apoptosis, and decreased cell division, thereby protecting against cancer cell proliferation (Hsieh et al., 2005; Takakuwa et al., 2019; Tucci, 2012). These findings suggest that in some group contexts, resource availability can be altered to promote either cooperation or selfishness. However, since cooperative and selfish entities compete at different levels of selection, both within and between groups, it is crucial to integrate the study of selection across these different levels to more fully understand how vital resources shape the competition dynamics between cooperative and selfish entities.

We sought to investigate the relationship between resource availability and multilevel selection on selfish and cooperative mitochondria. Cells normally contain dozens to thousands of mitochondrial organelles that undergo dynamic fusion and fission (Tilokani et al., 2018). These organelles in turn contain multiple copies of mitochondrial DNA (mtDNA), which are usually non-recombining and can replicate throughout the cell cycle (Chatre and Ricchetti, 2013; Newlon and Fangman, 1975; Sena et al., 1975). Mitochondria cooperate with each other and with their host by supplying energy; in return, the nuclear genome supplies the proteins and building blocks needed to replicate mtDNA. Selfish mtDNA can be defined as mutant variants that successfully propagate at the expense of host fitness (Taylor et al., 2002). Such mutants can coexist with their cooperative counterparts for many generations in a heteroplasmic state, hence multilevel selection shapes the population dynamics of mtDNA (Dubie et al., 2020; Klucnika and Ma, 2019; Rollins et al., 2016; Shou, 2015; Taylor et al., 2002). In yeast cells, for example, respiratory-deficient mutant mtDNA can outcompete wildtype mtDNA, resulting in the formation of “petite” colonies that fail to grow on non-fermentable food sources (Gaillard and Bernardi, 1979; Goldring et al., 1971; MacAlpine et al., 2001). In *Drosophila*, dysfunctional mtDNA copies can outcompete their functional counterparts within individual flies to the point of driving the fly stock extinct (Ma and O’Farrell, 2016). Some evidence suggests that selfish mtDNA might even persist in a heteroplasmic state over evolutionary timescales. For example, the same selfish mtDNA mutation has been identified in multiple geographically isolated strains of the nematode species *Caenorhabditis briggsae* (Howe and Denver, 2008).

How does resource availability influence the selection acting on selfish mtDNA? We address this question using a selfish mutant mtDNA within the model species *Caenorhabditis elegans*. First, we isolate and measure selection on the mutant mtDNA separately within individual hosts (sub-organismal) and between hosts (organismal). Consistent with the predictions of multilevel selection, sub-organismal selection favors the mutant mtDNA over the cooperative wildtype mtDNA, while the reverse is true at the level of organismal selection. Next, we applied this approach to study the effects of dietary nutrient availability and the physiology of metabolic stress tolerance. We show that although diet and nutrient sensing govern overall mtDNA levels, the preferential proliferation of ΔmtDNA at the sub-organismal level depends on the Forkhead box O (FoxO) transcription factor DAF-16. At the level of organismal selection, diet restriction promotes the relative fitness of the cooperative over the selfish mtDNA, but only in the absence of FoxO. This observation indicates that FoxO promotes host tolerance to the selfish mtDNA during food scarcity. Broadly speaking, these findings suggest that resilience to food scarcity influences the relative fitness of cooperators and cheaters both within and between groups.

## Results

### An experimental strategy to isolate selection on a selfish mitochondrial genome at different levels

Selection can act directly on individual mtDNA molecules within an organelle due to intrinsic replication advantage (Holt et al., 2014). Selection can also occur between organelles within a host cell (Lieber et al., 2019; Zhang et al., 2019), between cells within a multicellular host (Shidara et al., 2005), and finally between host organisms. The vast majority of mitochondrial content in the adult hermaphroditic nematode *Caenorhabditis elegans* is confined to the germline (Bratic et al., 2009), which exists as a contiguous syncytium of cytoplasm until the final stages of oocyte maturation (Pazdernik and Schedl, 2013). Sub-organismal selection thus predominantly reflects the biology of the female germline, where mtDNA variants compete for transmission to the next generation. Accordingly, here we focus on selection at the sub-organismal level as a single phenomenon, in addition to selection at the organismal level.

The present study takes advantage of the well-characterized heteroplasmic mutant genome *uaDf5* (Figure 1A), hereinafter referred to as ΔmtDNA (Ahier et al., 2018; Gitschlag et al., 2016; Liau et al., 2007; Lin et al., 2016; Tsang and Lemire, 2002). This deletion mutation spans four protein-coding genes and seven tRNA genes, disrupting gene expression and metabolic function. To quantify sub-organismal selection, we modified a previous approach to measure ΔmtDNA frequency in parents and their respective progeny as a function of parental ΔmtDNA frequency (Tsang and Lemire, 2002). Mutant frequency was measured longitudinally at successive developmental stages and across multiple parent-progeny lineages, which were propagated in isolation from one another to minimize the effect of organismal selection on overall ΔmtDNA frequency (Figures 1C-1D and S1). Initially, we observe reduced ΔmtDNA frequency in embryos compared to their parents (Figure 1B and 1C), consistent with germline purifying selection (Ahier et al., 2018; Hill et al., 2014; Lieber et al., 2019; Ma et al., 2014; Stewart et al., 2008). However, ΔmtDNA proliferates across development, achieving even higher frequency on average in adult progeny than in their respective parents (Figure 1C and 1D). Overall, we have quantitatively measured the frequency-dependent selection acting on ΔmtDNA at the sub-organismal level (Figure 1D).

**Figure 1.**
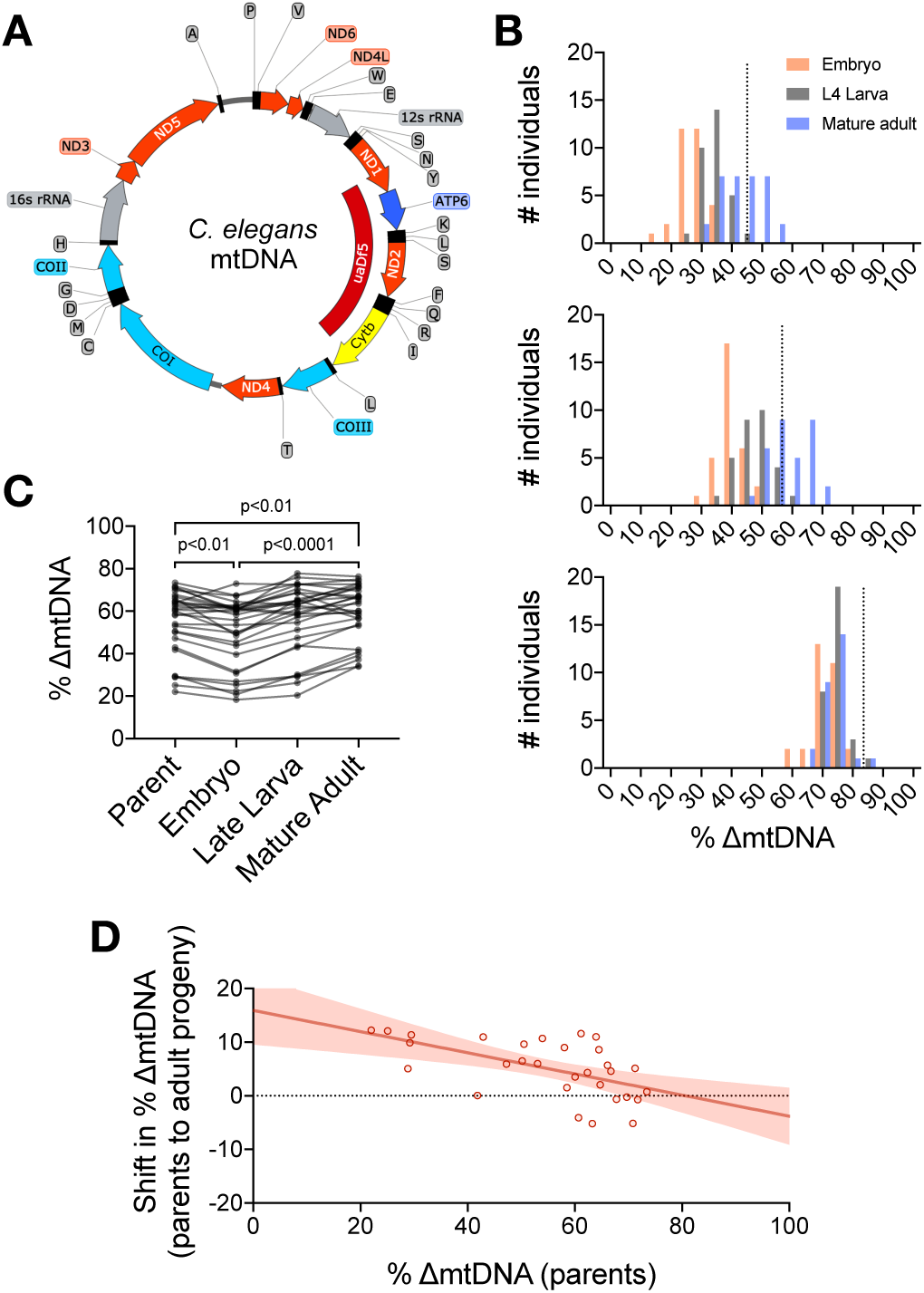
Quantification of sub-organismal selection for ΔmtDNA. (A) Map of *C. elegans* mtDNA showing the deletion that specifies the ΔmtDNA variant, *uaDf5* (dark red). Genes are color-coded according to functional category: respiratory complex I (light red), complex III (yellow), complex IV (light blue), complex V (dark blue), ribosomal RNA (gray), tRNA (black), non-coding regions (thin line). (B) Frequency distributions of ΔmtDNA across three life stages of single broods from parents with low (top, N=94), intermediate (middle, N=93), or high (bottom, N=88) ΔmtDNA frequency. Dotted lines represent parent ΔmtDNA frequency. (C) ΔmtDNA frequency between individual parents and age-synchronized progeny at three life stages. Each line is an individual parent-progeny lineage. Because parental ΔmtDNA frequency impacts fecundity and development (Figure 2C and 2D), individual parent-progeny lineages were propagated in isolation in order to minimize the effects of organismal fitness and magnify the effect of sub-organismal selection. Each parental data point represents a single parent lysed individually; each progeny data point represents three age-synchronized progeny lysed together to also minimize the effect of random drift. Friedman test with Dunn’s multiple comparisons test. (D) Shift in ΔmtDNA frequency per generation, between parents and their respective age-matched adult progeny. Plotted as a function of parental frequency using first and last time-points in (C). Red shaded region represents 95% C.I.

We observe several indicators that ΔmtDNA compromises host fitness (Figure 2A-2C), in agreement with previous studies of this genome (Gitschlag et al., 2016; Liau et al., 2007; Lin et al., 2016). To quantify selection against ΔmtDNA strictly at the level of host fitness, we competed heteroplasmic animals carrying ΔmtDNA against their homoplasmic wildtype counterparts on the same food plate (Figure 2D). In parallel, we propagated non-competing lines (lacking wildtype animals) to control for the confounding influence of sub-organismal heteroplasmy dynamics. By normalizing the population-wide changes in ΔmtDNA frequency to that of the non-competing control lines, we could measure selection against ΔmtDNA at the organismal level via the relative decline in ΔmtDNA frequency across the competing lines (Figures 2E and S2).

**Figure 2.**
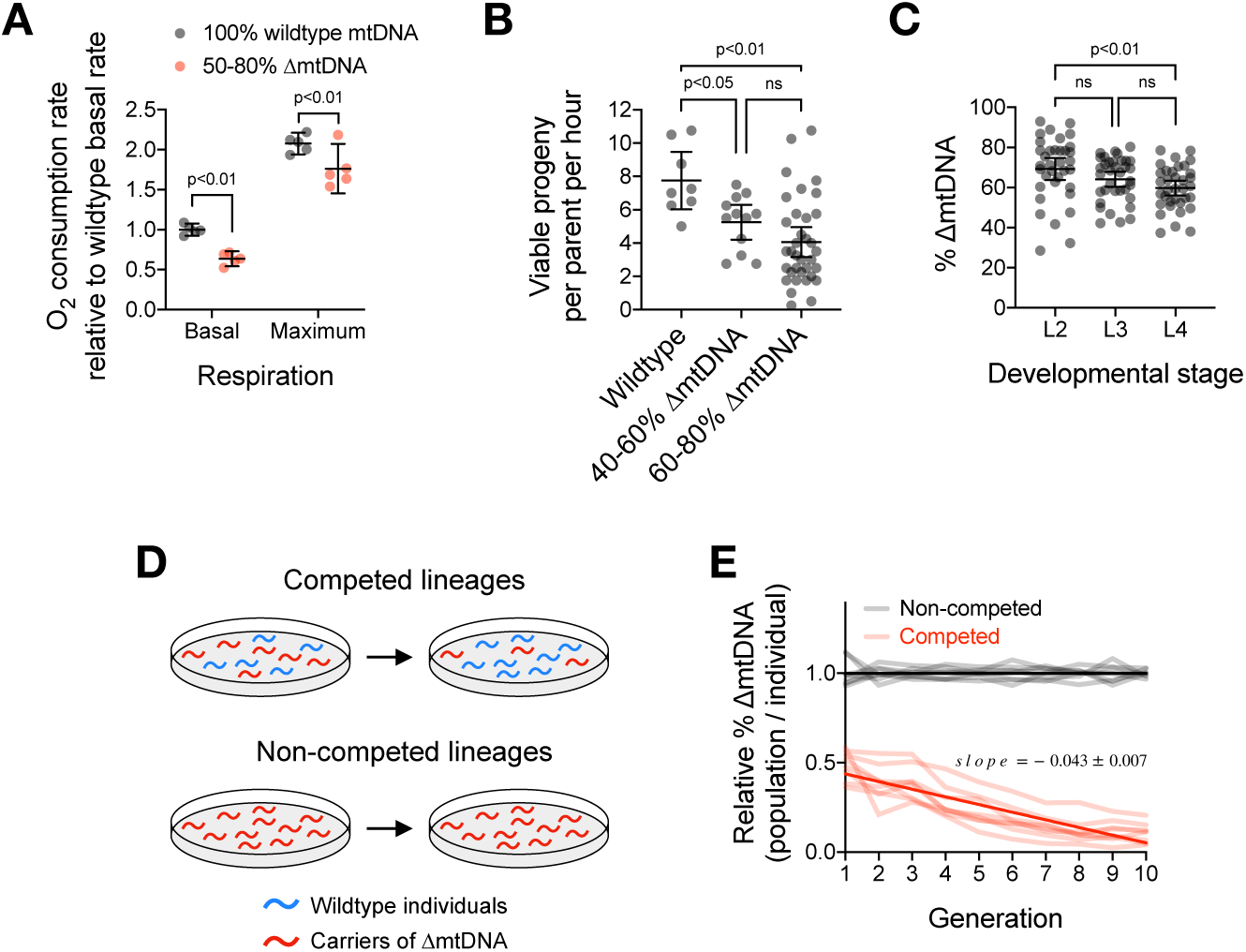
Quantification of organismal selection against ΔmtDNA. (A) Basal aerobic respiration and maximum respiratory capacity in age-synchronized adults maintaining ΔmtDNA. Two-way ANOVA with Sidak’s multiple comparisons test. (B) Peak fecundity as measured by viable progeny produced per hour from age-synchronized adults. Mitochondrial genotype was binned according to animals sampled from either the low end of the ΔmtDNA frequency distribution (defined as below the population mean of 60%, N=12), or the high end (defined as above the population mean of 60%, N=35), with wildtype controls (N=8). Brown-Forsythe and Welch ANOVA with Dunnett’s T3 multiple comparisons test. (C) ΔmtDNA frequency between larval stages among a chronologically synchronized cohort. N=35 per developmental stage. Brown-Forsythe and Welch ANOVA with Dunnett’s T3 multiple comparisons test. Each data point represents one nematode. (D) Schematic illustrating experimental design for competition experiment to quantify organismal selection against ΔmtDNA. Heteroplasmic carriers of ΔmtDNA used in competing and non-competing lineages were taken from the same stock. (E) Population-wide ΔmtDNA frequency, relative to ΔmtDNA frequency per heteroplasmic individual, measured for 10 generations across 8 replicate competed (red) and non-competed (gray) lineages. Competed lineages consisted of ΔmtDNA-carrying heteroplasmic individuals mixed with homoplasmic wildtype counterparts on the same food plates. Non-competed lineages consisted of exclusively ΔmtDNA-carrying heteroplasmic individuals. Lower initial population-wide ΔmtDNA frequency in competing lines is due to the presence of wildtype animals. At each generation, population-wide ΔmtDNA frequency of competing and non-competing lines were sampled and normalized to average population-wide frequency of all non-competing lines (which is equivalent to average frequency per heteroplasmic individual, since all individuals within non-competing lines carry ΔmtDNA). Solid lines represent best-fit regressions across all replicate lineages. Error bars represent 95% C.I.

We sought to integrate these measurements of organismal and sub-organismal selection into a single mathematical framework to describe the overall population dynamics of ΔmtDNA. The Price equation provides such a framework by describing evolution in terms of the covariance between a measurable trait value and its associated reproductive fitness (Frank, 1997; Price, 1972). Since the Price equation can accommodate more than one source of covariance, it is ideal for characterizing selection at multiple levels:

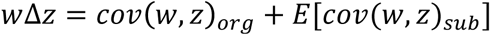

Here, *z* refers to a character trait, in this case mitochondrial genotype. Since mitochondrial genotype consists of a heteroplasmic mix of ΔmtDNA and wildtype mtDNA, we define the trait value *z* as the heteroplasmic frequency of ΔmtDNA, while *w* refers to the fitness or reproductive rate of ΔmtDNA relative to that of wildtype mtDNA. The subscripts *_org_* and *_sub_* label the covariance between *w* and *z* at the organismal and sub-organismal levels, respectively. The expression *E* refers to expected, or average, sub-organismal covariance between *w* and *z*. Moreover, the strength with which natural selection acts on a genotype can be expressed as the selection coefficient, *s* (equal to 1–*w*). Hence, the change in a trait due to selection can be described using the covariance between trait value and selection coefficient. Summing the organismal and sub-organismal covariances then provides a description of the overall population-level selection dynamics of ΔmtDNA.

We converted the measured sub-organismal shifts in ΔmtDNA frequency per generation (Figure 1D) to selection coefficients and plotted them as a function of parental ΔmtDNA frequency (Figure 3, top graph). At a heteroplasmic ΔmtDNA frequency of approximately 60%, we found that the organismal selection coefficient for selection against ΔmtDNA is 0.23±0.04 (95% C.I.) across 8 replicate lineages (Figures 2E and S2A). The organismal selection coefficient would be zero when ΔmtDNA frequency is 0%, and the selection coefficient would be 1.0 when ΔmtDNA frequency is 100% since the mutation deletes essential genes and is thus predicted to be lethal at 100%. Based on these data points, we derived a non-linear regression between ΔmtDNA frequency and selection coefficient (Figure 3, middle graph). This regression predicts that the organismal fitness cost accelerates with increasing ΔmtDNA frequency, which is consistent with the notion of a phenotypic threshold effect as mutant mtDNA levels rise (Letellier et al., 1994; Picard et al., 2014; Sciacco et al., 1994). Summing the sub-organismal and organismal regressions yields the overall population-level relationship between ΔmtDNA frequency and selection (plotted in Figure 3, bottom graph):

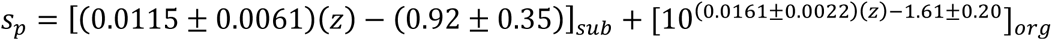

**Figure 3.**
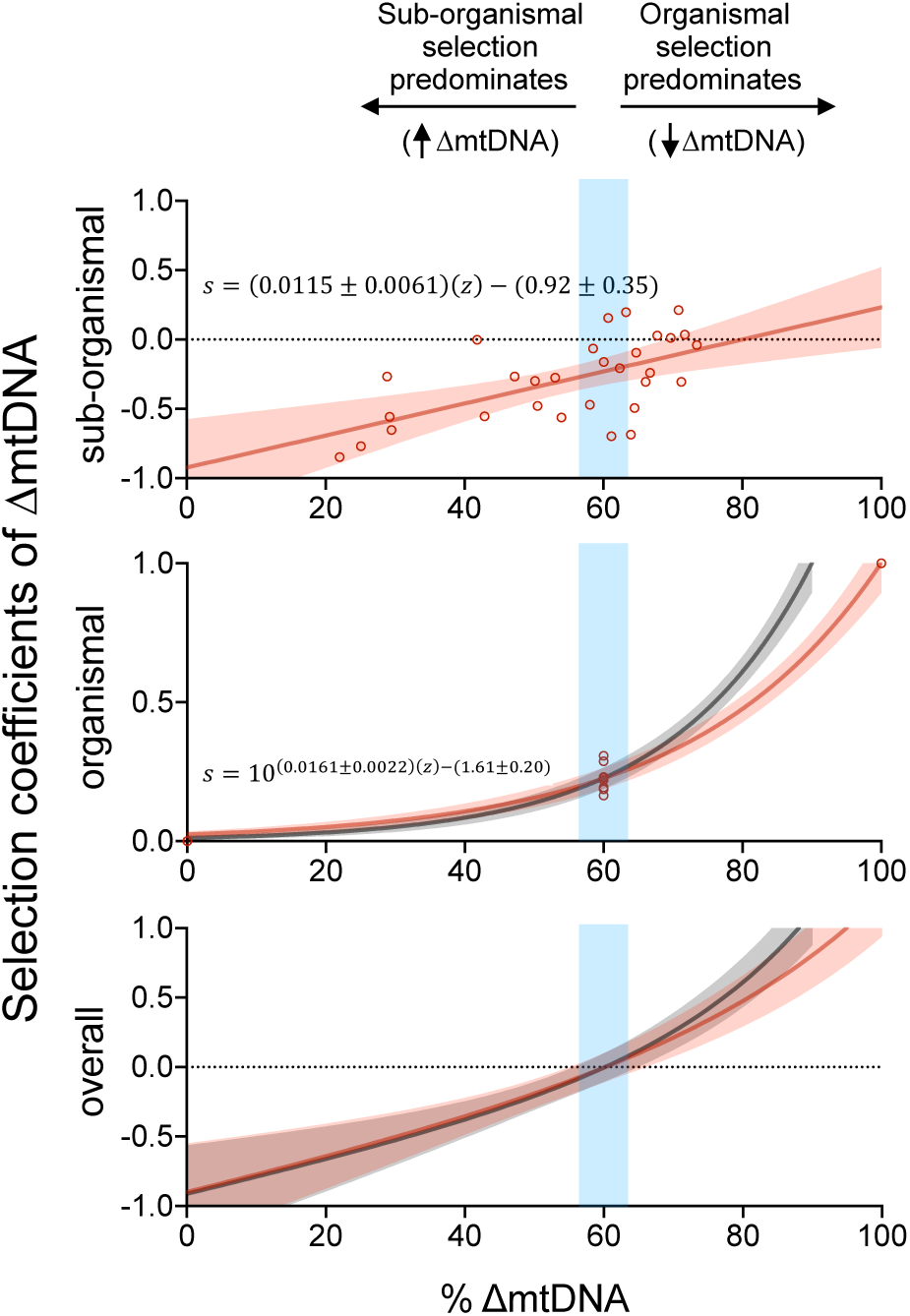
Fitness of wildtype mtDNA relative to ΔmtDNA as a function of ΔmtDNA frequency, at each selection level. Selection coefficient against ΔmtDNA at sub-organismal (top), organismal (middle), and overall combined (bottom) levels of selection. The top graph represents the same regression as in Figure 1D but with absolute ΔmtDNA frequency shift converted to selection coefficient. When selection coefficient against ΔmtDNA is negative, positive selection is acting on ΔmtDNA, hence frequency rises. At 100% ΔmtDNA frequency, the mutation is lethal and hence the selection coefficient against ΔmtDNA is expected to be 1.0 (middle graph, upper right corner). At 0% frequency, no ΔmtDNA is available for selection, hence the organismal selection coefficient against ΔmtDNA is 0. At 60% ΔmtDNA frequency, the empirically derived selection coefficient against ΔmtDNA is 0.23 (95% C.I. ±0.04). The plotted curve represents the line that passes through all three of these values for selection coefficient. The blue shaded region represents the range of ΔmtDNA frequency in which the organismal and sub-organismal selection coefficients balance, resulting in a stably persisting ΔmtDNA near 60% frequency. Red regions represent 95% C.I. Gray shaded regions (middle and lower graphs) show regressions when selection coefficient reaches 1.0 at 90% ΔmtDNA frequency.

Here, *s_p_* represents the overall (population-level) selection coefficient against ΔmtDNA while the subscripts *_sub_* and *_org_* label the sub-organismal and organismal regressions, respectively.

To derive this model, we assumed that ΔmtDNA becomes non-viable upon reaching 100% frequency. However, ΔmtDNA is rarely observed above 90% frequency (Gitschlag et al., 2016; Liau et al., 2007). To account for the uncertainty in how organismal selection changes with rising ΔmtDNA frequency, we set the organismal selection coefficient to 1.0 when ΔmtDNA frequency reaches 90% (Figure 3, gray shaded region). We also tested the assumption that organismal selection increases linearly with ΔmtDNA frequency (Figure S3). These alternate models had negligible effect on our overall conclusions. In each case, positive sub-organismal selection and negative organismal selection balance when ΔmtDNA is near 60% mean heteroplasmic frequency. In summary, we have separately measured selection on ΔmtDNA at each level and integrated them into the mathematical framework of the Price equation, which robustly predicts the population-level dynamics of ΔmtDNA that we observe.

### ΔmtDNA exploits nutrient status to propagate

Using this framework, we sought to investigate how resource availability affects the multilevel selection acting on ΔmtDNA. Given the central role of mitochondria in metabolism, we reasoned that the multilevel selection acting on ΔmtDNA might be particularly sensitive to dietary perturbations. We therefore raised nematodes on food plates seeded with either a high or low concentration of *E. coli* (OP50 strain), which were UV-killed to prevent further bacterial growth (Figure S5A-S5B). UV-killed OP50 partially mimics diet restriction (Win et al., 2013). Nevertheless, we found that nematodes raised on the more restricted (low concentration) diet during the transition from larvae to mature adulthood—when the greatest net increase in mtDNA copy number occurs—harbored significantly lower ΔmtDNA frequency compared to those raised on a more abundant control (high concentration) diet (Figure 4A). Moreover, we observed that sub-organismal ΔmtDNA frequency is even higher when animals are raised on live food (Figure S5C). Tracking mutant frequency across development revealed that ΔmtDNA levels rise from embryos to adults on a control diet as expected, but not in animals grown on a restricted diet (Figure 4B and 4C). Based on these observations, we conclude that ΔmtDNA exploits nutrient abundance to rise in frequency during development but fails to do so under conditions of diet restriction.

**Figure 4.**
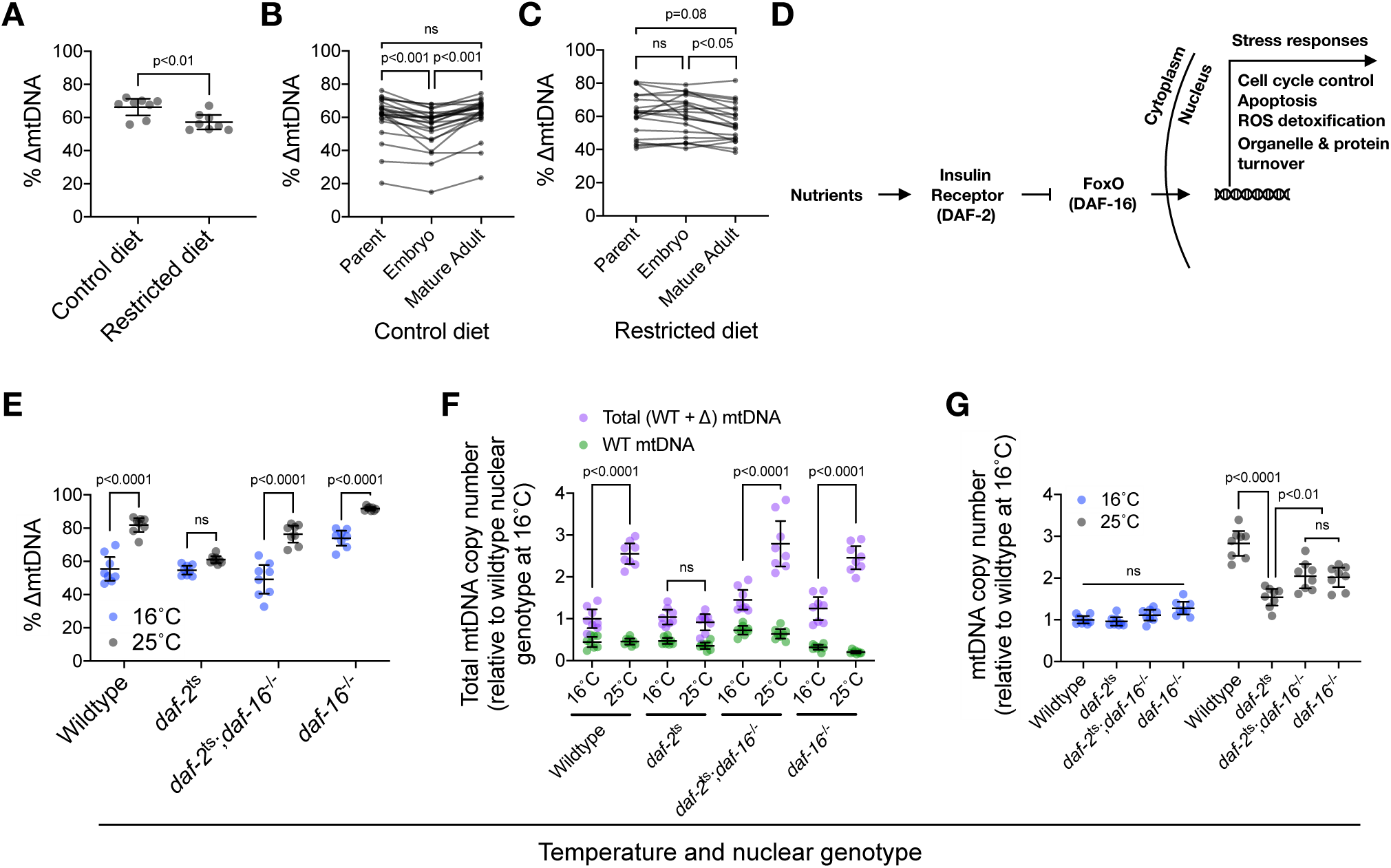
ΔmtDNA exploits nutrient supply and insulin signaling to proliferate at the sub-organismal level. (A) Frequency of ΔmtDNA between age-synchronized adults raised on a restricted versus control diet. Mann-Whitney test. N=8 lysates containing 5 pooled age-synchronized adults per lysate. (B-C) ΔmtDNA levels in individual parents and age-synchronized progeny at two life stages, embryo and mature adult, across lineages raised on either control (B) or restricted (C) diet. Each parental data point represents a single parent lysed individually; each progeny data point represents three age-synchronized progeny lysed together. Friedman test with Dunn’s multiple comparisons test. N=24 parent-progeny lineages (control diet); N=20 parent-progeny lineages (restricted diet). (D) Schematic of the FoxO-dependent insulin signaling cascade, with *C. elegans* homologs of mammalian proteins in parentheses. (E) Frequency of ΔmtDNA in age-synchronized adults of wildtype, temperature-sensitive *daf-2*(e1370) mutant, null *daf-16*(mu86) mutant, or double-mutant genotype. Nematodes were raised from late larval (L4) to mature adulthood at 25°C, the restrictive temperature for *daf-2*(e1370), with controls raised at the permissive temperature of 16°C. Two-way ANOVA with Bonferroni correction. N=8 lysates containing 5 pooled age-synchronized adults each. (F) mtDNA copy number of the same samples shown in (E), separated by mitochondrial genotype and normalized to total mtDNA of adults with the wildtype nuclear genotype raised at the control temperature of 16°C. Wildtype mtDNA copy number is shown in green, total mtDNA copy number is shown in purple. The vertical distance between wildtype and total mtDNA represents ΔmtDNA copy number. Two-way ANOVA with Bonferroni correction. (G) mtDNA copy number in homoplasmic age-synchronized adults of wildtype, temperature-sensitive *daf-2*(e1370) mutant, null *daf-16*(mu86) mutant, or double-mutant genotype, raised from L4 stage at the restrictive temperature of 25°C, with controls maintained at the permissive temperature of 16°C. Two-way ANOVA with Bonferroni correction. N=8 lysates containing 5 pooled age-synchronized adults each. Error bars represent 95% C.I.

### Insulin signaling influences sub-organismal proliferation of ΔmtDNA

To better understand the role of nutrient availability in sub-organismal ΔmtDNA proliferation, we targeted the insulin-signaling pathway. Insulin acts as a nutrient-dependent growth hormone and regulator of metabolic homeostasis, tailoring the appropriate physiological responses to external nutrient conditions (Badisco et al., 2013; Danielsen et al., 2013; Das and Arur, 2017; Lee and Dong, 2017; Lopez et al., 2013; Michaelson et al., 2010; Porte et al., 2005; Puig and Tjian, 2006; Shiojima et al., 2002). Nematodes expressing a defective allele of the insulin receptor homolog *daf-2* perceive starvation, even in presence of food. However, disrupting insulin signaling in young larvae causes the dauer phenotype, a form of developmental arrest (Gottlieb and Ruvkun, 1994). Thus, we cannot directly track ΔmtDNA frequency across development under prolonged loss of *daf-2* function. Instead, we used a temperature-sensitive *daf-2* allele to bypass dauer arrest. We incubated these animals at a permissive temperature (16°C) during early larval development, preserving insulin signaling. Next, we shifted these animals to the restrictive temperature (25°C) at a later larval (L4) stage, thereby disrupting insulin signaling during the transition from larva to mature adult. In control animals expressing a wildtype receptor, adult maturation at 25°C coincides with rising ΔmtDNA frequency compared to maturation at 16°C (Figure 4E). However, this dramatic increase in ΔmtDNA frequency is abolished in temperature-sensitive *daf-2* mutants (Figure 4E). Indeed, we observed no overall change in average ΔmtDNA frequency across four independent lineages of the temperature-sensitive *daf-*2 mutants even after four consecutive generations of adult maturation at 25°C and subsequent recovery of larval progeny at 16°C (Figure S5D). In contrast, we observed a robust increase in ΔmtDNA frequency in wildtype controls. We therefore conclude that nutrient sensing via the insulin-signaling pathway regulates the sub-organismal proliferation of ΔmtDNA.

The insulin receptor communicates nutrient status to the cell largely through the negative regulation of the FoxO family of transcription factors (O-Sullivan et al., 2015), encoded by the gene *daf-16* in *C. elegans* (Figure 4D). Nutrient limitation or inactivation of the receptor activates FoxO/DAF-16, resulting in altered expression of its target genes. We therefore hypothesized that DAF-16 functions downstream of DAF-2 to suppress ΔmtDNA proliferation. Indeed, deletion of *daf-16* restores the temperature-dependent proliferation of ΔmtDNA in animals expressing the temperature-sensitive *daf-2* allele (Figure 4E). These data support a role for DAF-16 in regulating ΔmtDNA proliferation.

Next, we sought to understand how DAF-2-dependent inhibition of DAF-16 regulates ΔmtDNA proliferation. In addition to ΔmtDNA frequency, we also found that total mtDNA copy number rises at 25°C compared to 16°C (Figure 4F). Interestingly, increased copy number of ΔmtDNA entirely accounts for the rise in total copy number. Furthermore, higher total copy number at 25°C was abolished in *daf-2* mutants and restored in *daf-2*;*daf-16* double-mutants (Figure 4F). To understand whether rising total mtDNA copy number promotes ΔmtDNA proliferation, or merely results from it, we quantified copy number across the same conditions in animals lacking ΔmtDNA. Homoplasmic-wildtype mtDNA copy number rose at 25°C compared to 16°C (Figure 4G). This rise in copy number was suppressed in *daf-2* mutants and partially but significantly rescued in *daf-2*;*daf-16* double-mutants (Figure 4G). Moreover, RNAi knockdown of *daf-2* expression also suppressed mtDNA copy number in a *daf-16*-dependent manner (Figure S5E). Together, these data show that DAF-2 signaling inhibits DAF-16 to allow high mtDNA copy number, which permits sub-organismal ΔmtDNA proliferation at the warmer temperature.

How might DAF-16 suppress mtDNA copy number upon loss of DAF-2 signaling? DAF-16 could achieve copy-number suppression via elimination of mitochondria, either through mitochondrial fragmentation and subsequent mitochondrial autophagy, or at the cellular level through apoptosis. However, these processes do not appear to play a role in mtDNA copy number suppression in *daf-2* mutants (Figures 5A-5C and S6). Alternatively, DAF-16 might suppress mtDNA copy number by limiting mitochondrial biogenesis. Nutrient availability and insulin signaling each promote development of the germline (Angelo and Van Gilst, 2009; Drummond-Barbosa and Spradling, 2001; Michaelson et al., 2010; Narbonne and Roy, 2006; Shim et al., 2002), which harbors the vast majority of mtDNA in the adult nematode (Bratic et al., 2009). We observed that mitochondrial organelle quantity and mtDNA copy number are proportional to gonad size across wildtype, *daf-2* mutant, and *daf-2*;*daf-16* double-mutants (Figure 5D-5G). These data indicate that upon loss of insulin signaling, DAF-16 suppresses germline development and the accompanying biogenesis of mtDNA.

**Figure 5.**
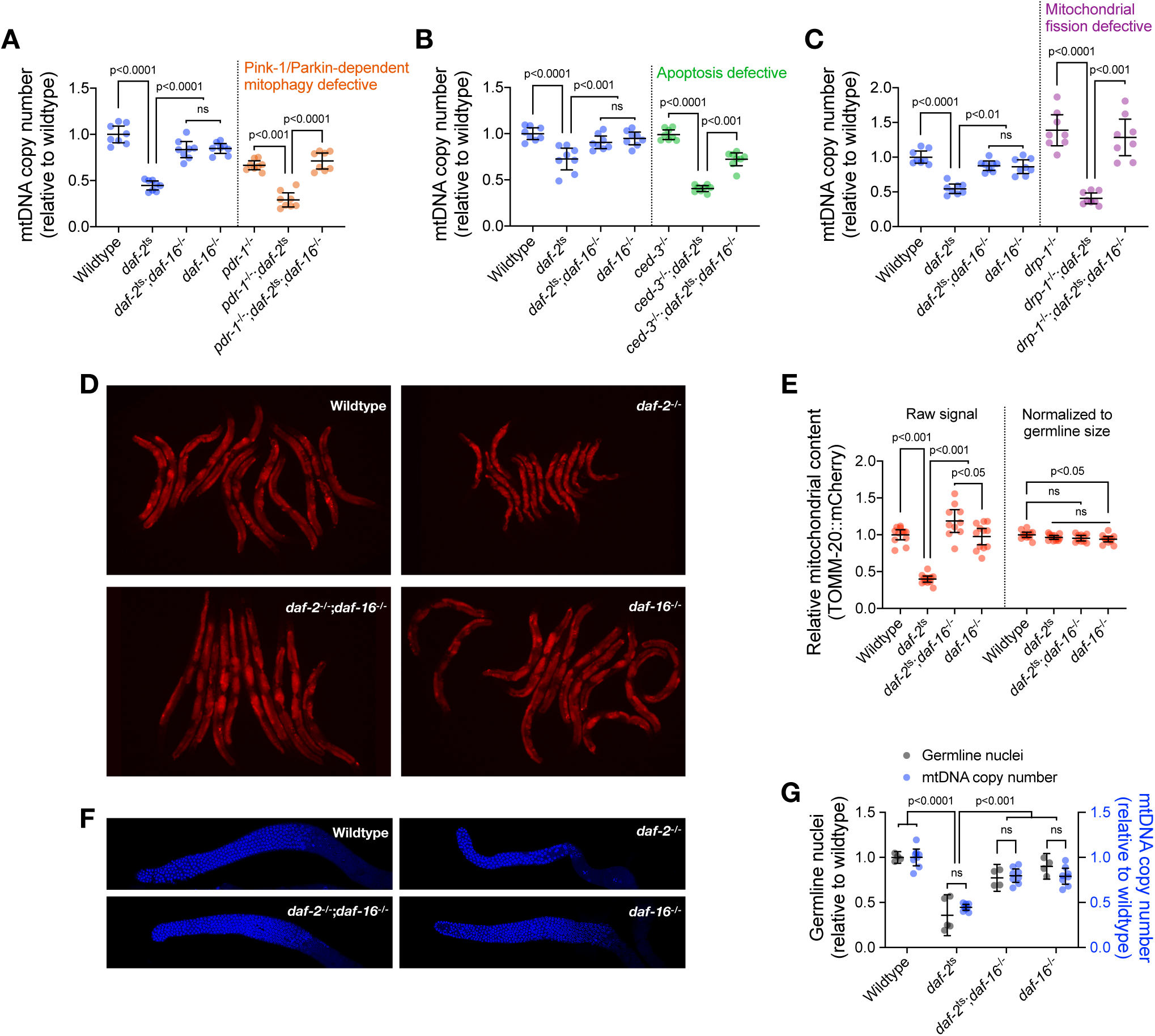
DAF-16 activation upon loss of insulin signaling suppresses mtDNA content via regulation of germline proliferation. (A-C) mtDNA copy number in age-synchronized adults of wildtype, temperature-sensitive *daf-2*(e1370) mutant, null *daf-16*(mu86) mutant, or double-mutant genotype, each raised from L4 stage at 25°C. Copy number is also shown in wildtype, *daf-2*(e1370), and *daf-2*(e1370);*daf-16*(mu86) double-mutant adults each with *pdr-1*(gk448) (A), *ced-3*(ok2734) (B), or *drp-1*(tm1108) (C), representing loss-of-function alleles of the Parkin homologue, a terminator caspase, and dynamin-related protein, respectively. mtDNA copy number in *daf-16*(mu86) single-mutants is also shown. One-way ANOVA with Bonferroni correction. N=8 lysates containing 5 pooled age-synchronized adults each. (D-E) Images and quantification of germline mitochondrial organelle signal intensity as indicated by the fluorescent reporter TOMM-20::mCherry, across age-synchronized adults of wildtype, temperature-sensitive *daf-2*(e1370) mutant, null *daf-16*(mu86) mutant, or double-mutant genotype, raised from L4 stage at 25°C. Each data point in (E) represents one age-synchronized adult nematode, shown in (D). One-way ANOVA with Bonferroni correction. (F-G) Images and quantification of DAPI-stained nuclei and mtDNA copy number of age-synchronized adults of wildtype, temperature-sensitive *daf-2*(e1370) mutant, null *daf-16*(mu86) mutant, or double-mutant genotype, raised from L4 stage at 25°C. Each nuclei data point in (G) represents one female gonad (age-synchronized adults); each mtDNA copy number data point represents a lysate containing 5 pooled age-synchronized adults. Two-way ANOVA with Bonferroni correction. Error bars represent 95% C.I.

The observation that DAF-16 suppresses mtDNA copy number in insulin-signaling mutants suggests that DAF-16 may also suppress mtDNA copy number in response to diet restriction. However, total mtDNA copy number was substantially lower in heteroplasmic animals raised on the restricted diet compared to the control diet, even in *daf-16* mutants (Figure 6A). These data show that diet restriction limits copy number independently of DAF-16. Given that elevated copy number is associated with higher ΔmtDNA frequency (Figure 4E and 4F), ΔmtDNA frequency may also benefit from dietary nutrient abundance independently of DAF-16. Remarkably, ΔmtDNA frequency was higher on the more plentiful control diet compared to the restricted diet, but only when DAF-16 was present (Figure 6B). Moreover, while total mtDNA copy number and ΔmtDNA frequency each rose significantly across development on the control diet compared to the restricted diet (Figure 6C), copy number increased without any accompanying change in ΔmtDNA frequency in *daf-16* mutants (Figure 6D). Together, these data reveal two opposing roles for DAF-16. Upon loss of insulin signaling, DAF-16 suppresses germline development and mtDNA copy number, limiting the niche space in which ΔmtDNA can proliferate. Conversely, DAF-16 is required for preferential ΔmtDNA proliferation under conditions of nutrient abundance. We therefore conclude that nutrient abundance and DAF-16 are each necessary, but not sufficient individually, for the sub-organismal selection advantage of ΔmtDNA.

**Figure 6.**
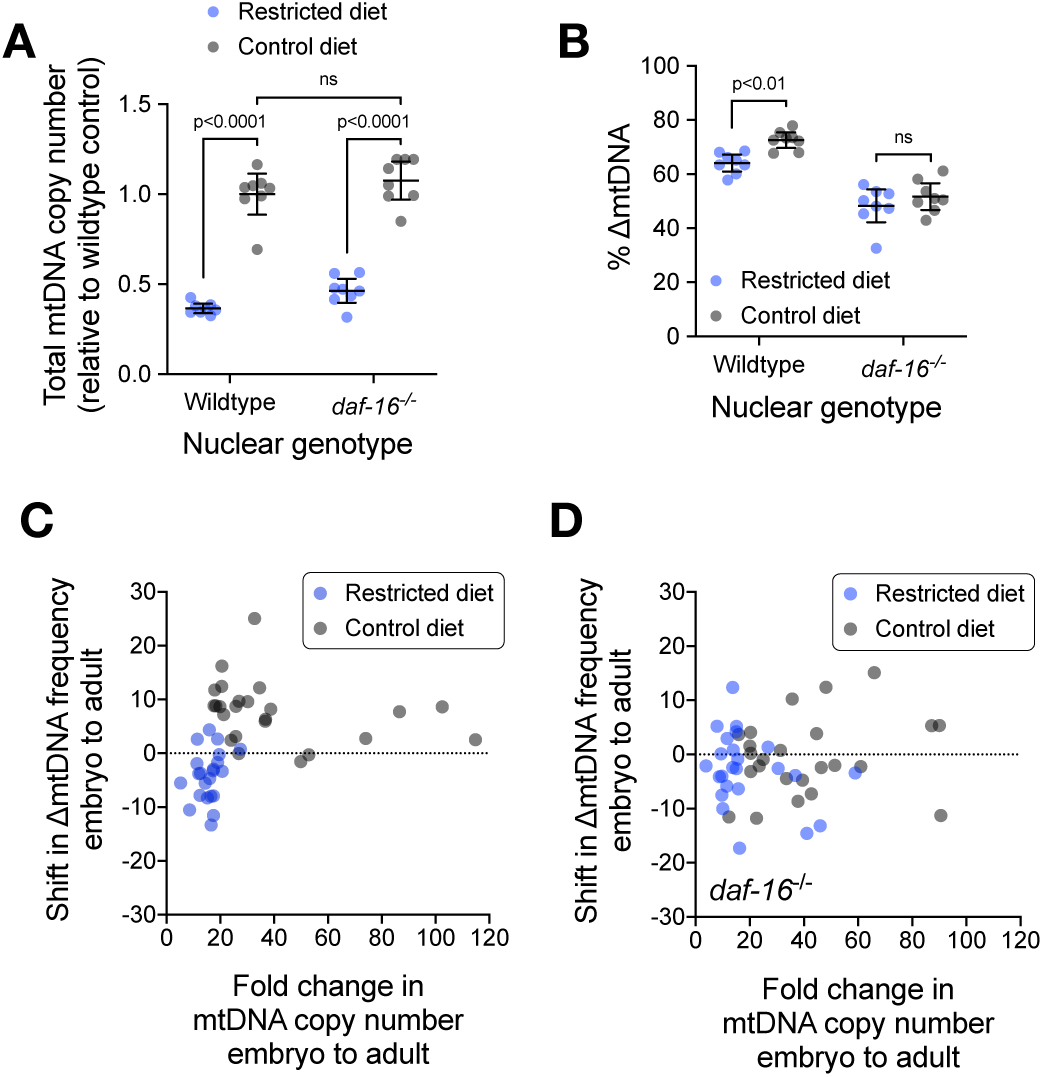
The sub-organismal selection advantage of ΔmtDNA requires both nutrient abundance and DAF-16. (A) Total mtDNA copy number across heteroplasmic age-synchronized adults of wildtype nuclear genotype or null *daf-16*(mu86) genotype, raised on a restricted diet versus a control diet. Two-way ANOVA with Bonferroni correction. N=8 lysates containing 5 pooled age-synchronized adults each. (B) ΔmtDNA frequency across the samples shown in (A). Two-way ANOVA with Bonferroni correction. (C-D) Shift in ΔmtDNA frequency across development as a function of change in mtDNA copy number, in control and restricted diets, across animals of wildtype nuclear genotype (C) or null *daf-16*(mu86) genotype (D). Each data point corresponds to an embryonic lysate (3 pooled embryos) and a corresponding adult lysate (3 pooled adults) taken from the same parent and lysed 96 hours apart. Mann-Whitney tests with Bonferroni correction. N=22 (restricted diet, wildtype); N=24 (control diet, wildtype); N=24 (restricted diet, *daf-16*^-/-^); N=24 (control diet, *daf-16*^-/-^). Error bars represent 95% C.I.

### Nutrient status governs selection on ΔmtDNA at different levels

Dietary nutrient abundance and DAF-16 each affect sub-organismal ΔmtDNA frequency (Figure 6A-6D), and ΔmtDNA frequency itself affects host fitness. Additionally, FoxO/DAF-16 regulates numerous genes involved in stress tolerance (Klotz et al., 2015; Martins et al., 2016; Murphy et al., 2003; Tepper et al., 2013; Webb et al., 2016) and promotes organismal survival during nutrient scarcity (Greer et al., 2007; Hibshman et al., 2017; Kramer et al., 2008). Given these considerations, we hypothesized that dietary nutrient availability and DAF-16 affect selection on ΔmtDNA at both the organismal and sub-organismal levels. Sub-organismal selection was quantified as before (see Figure 1D), under restricted versus control diets, in the presence versus absence of DAF-16 (Figure 7A and 7B). Organismal selection was quantified under each of these same conditions, using the competition method previously described (see Figure 2D and 2E). In populations with wildtype DAF-16, diet restriction did not significantly affect the decline in ΔmtDNA frequency at the level of organismal selection (Figures 7C, S7A and S7D). However, diet restriction accelerated the selection against ΔmtDNA at the organismal level in competing populations of *daf-16* mutants (Figures 7D, S7B, S7C and S7E). These data indicate that on a restricted diet, DAF-16 helps curtail organismal selection against ΔmtDNA.

**Figure 7.**
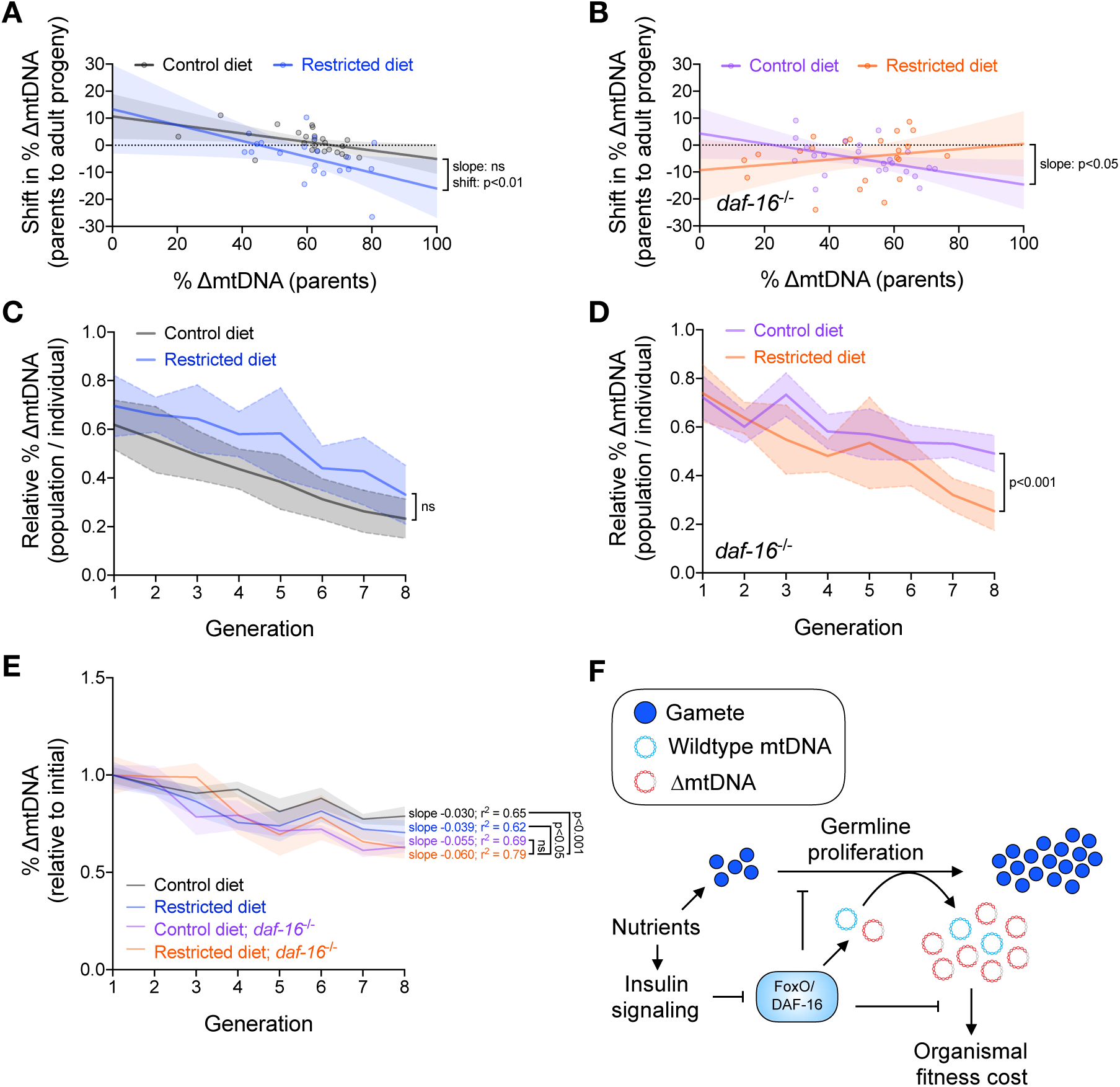
Nutrient status impacts multilevel selection dynamics of ΔmtDNA. (A-B) Shift in ΔmtDNA frequency per generation attributable to sub-organismal mitochondrial dynamics, similar to Figure 1D, featuring parent-progeny lineages raised on a restricted or control diet, across animals of wildtype nuclear genotype (A) or null *daf-16*(mu86) genotype (B). Linear regression compares parental ΔmtDNA frequency with the magnitude of frequency-shift from parent to age-matched progeny. Regressions compared using analysis of covariance. (C-D) Measure of organismal selection against ΔmtDNA, similar to Figure 2B, in competing lineages raised on a restricted or control diet, across animals of wildtype nuclear genotype (C) or null *daf-16*(mu86) genotype (D). Vertical axis represents relative ΔmtDNA frequency in the population, measured as proportion of average ΔmtDNA frequency per heteroplasmic individual (equal to population-wide ΔmtDNA frequency of non-competing populations). (E) ΔmtDNA frequency, relative to starting frequency, in non-competing lineages from the organismal competition experiment shown in (C) and (D). Linear regression analyses with Bonferroni correction. (F) Model illustrating the influence of DAF-16 on ΔmtDNA selection dynamics. Dietary nutrient supply and the inhibition of DAF-16 by insulin signaling promote germline growth and mtDNA replication (Figures 4 and 5). However, the ability of ΔmtDNA to take advantage of nutrient supply and preferentially proliferate requires DAF-16 (Figure 6), indicating that nutrient supply and DAF-16 are each necessary, but not sufficient individually, for ΔmtDNA proliferation. Organismal selection against ΔmtDNA intensifies during dietary restriction but only in the absence of DAF-16, indicating that DAF-16 partially shields ΔmtDNA from organismal selection during conditions of food scarcity. Hence, nutrient status and host genotype interact to shape the fitness of ΔmtDNA across the levels of selection.

Finally, we used the framework of the Price equation to integrate the sub-organismal and organismal covariances for each of the four conditions tested (Figure S7D and S7E). The combined covariances predict that the strongest overall selection against ΔmtDNA occurs in populations lacking DAF-16 experiencing food scarcity, and the weakest overall selection occurs in populations with DAF-16 experiencing food abundance, with the remaining two conditions each experiencing an intermediate strength of selection (Figure S7D and S7E, bottom graphs). Measuring ΔmtDNA frequency across non-competing heteroplasmic populations afforded the opportunity to test this prediction. Remarkably, this prediction is consistent with our observation (Figure 7E). Combined, our data show that quantitatively accounting for selection at each level allows for a fuller description of the population dynamics of a cheater genome, including the effects of food scarcity and host genotype on those dynamics (Figure 7F).

## Discussion

Deleterious fitness effects are no guarantee that natural selection will remove a trait from the population. On the contrary, traits that are deleterious at one level of scale can be favored at a different level, a phenomenon known to characterize mtDNA heteroplasmy dynamics (Clark et al., 2012; Dubie et al., 2020; Gitschlag et al., 2016; Liau et al., 2007; Lin et al., 2016; Ma and O’Farrell, 2016; Taylor et al., 2002; Tsang and Lemire, 2002). By isolating and measuring selection at each level, we have applied a theoretical framework to empirical data to uncover the impact of environmental context on multilevel selection. Using this approach, we identified a diet-by-genotype interaction that influences the population genetics of mitochondria in *C. elegans*.

We have shown here that although dietary nutrient abundance fuels mtDNA replication, the preferential sub-organismal proliferation of ΔmtDNA depends on DAF-16 and its downstream functions. Our study provides generalizable insights into conditions that determine the fitness of cheaters versus cooperators. In particular, we find that the opportunity for cheaters to outcompete cooperators requires resources and a niche space where replication can occur. However, while necessary, these conditions are not sufficient to ensure the fitness advantage of cheaters. On the contrary, additional conditions shape the cost of cooperation, or the benefit of cheating, and hence the fitness advantage of cheaters.

How does DAF-16 promote ΔmtDNA proliferation? This could occur via several potential mechanisms. For example, a recent study in heteroplasmic flies found that PINK1 preferentially localizes to mitochondria enriched for the mutant genome in ovaries, inhibiting protein synthesis and mtDNA replication (Zhang et al., 2019). Furthermore, previous genome-wide expression analysis identified DAF-16 as a potential negative regulator of the gene encoding PINK1 (Tepper et al., 2013), suggesting that DAF-16 might interfere with the PINK1-dependent preferential replication of wildtype mtDNA, although it is not known whether this function of PINK1 is evolutionarily conserved between flies and nematodes. Another recent study from the same group found that insulin receptor signaling functions through the protein Myc to limit the propagation of mutant mtDNA in heteroplasmic fly ovaries (Wang et al., 2019). Interestingly, Myc has also been identified among the target genes of FoxO transcription factors in the fly soma (Teleman et al., 2008) as well as in mammals (Webb et al., 2016), suggesting that DAF-16 might function through a similar mechanism to regulate heteroplasmy dynamics in nematodes. Alternatively, DAF-16 could promote ΔmtDNA proliferation via the alleviation of bioenergetic stress. DAF-16 up-regulates the expression of multiple genes involved in energy metabolism, including several glycolytic enzymes (Depuydt et al., 2014; Tepper et al., 2013). By promoting the mitochondria-independent production of ATP, DAF-16 might be contributing to metabolic conditions that make the cell more tolerant to the presence of mutant mitochondria. This possibility is consistent with previous work implicating other compensatory stress-response mechanisms in the proliferation of ΔmtDNA (Gitschlag et al., 2016; Lin et al., 2016).

How does resource availability influence selection on cooperators versus cheaters at the level of competing groups? Note that the female germline, along with the cell lineage that develops from an oocyte to a germline in the next generation, harbors the mtDNA molecules that compete for transmission. Organismal selection acting on mtDNA can thus be viewed as a group-level phenomenon, while sub-organismal selection represents the within-group level. On one hand, if resource scarcity selects for cooperation, then groups with a higher proportion of cooperators should experience an additional fitness advantage over other groups during times of scarcity. On the other hand, exposure to cheating can result in an evolutionary arms race in which cooperators acquire resistance to cheaters, a phenomenon observed in bacteria and social amoebae (Hollis, 2012; Khare et al., 2009; O’Brien et al., 2017). Could food scarcity select for adaptations that reduce the fitness impact of metabolic cheaters? We propose that DAF-16 represents an example of this type of stress tolerance. Although diet restriction compromised the sub-organismal advantage of ΔmtDNA, it had no effect on the organismal disadvantage, provided DAF-16 is present. However, in *daf-16* mutants, diet restriction intensified organismal selection against ΔmtDNA. FoxO/DAF-16 is known to promote survival under conditions of nutrient limitation (Greer et al., 2007; Hibshman et al., 2017; Kramer et al., 2008). Here, we have shown that the same gene also promotes host tolerance to the presence of ΔmtDNA during food scarcity.

It is important to note that we do not draw specific inferences about the genetic composition of naturally occurring populations. For example, this study does not assume that nuclear or mitochondrial mutants would necessarily self-segregate into separate populations in the wild. Likewise, organisms carrying a deleterious heteroplasmy are unlikely to spread through a population except in rare circumstances, such as following an extreme population bottleneck. Rather, we sought to uncover basic principles describing how resource availability and multilevel selection interact to shape the population dynamics of cooperative and selfish entities. In conclusion, the work presented here suggests that when cooperators and cheaters compete across different levels of selection, resource availability—and resilience to resource scarcity—together shape the proliferative capacity of the cheaters, as well as the ability of the cooperators to collectively tolerate their presence.

## Acknowledgments

We thank the members of the Patel Laboratory (James P Held, Cait S Kirby, Nikita Tsyba, Benjamin Saunders, Cassidy A Johnson), Janet M Young, Mia T Levine, Sarah E Zanders, Harmit S Malik, and Antonis Rokas for their valuable feedback on the manuscript. This work was generously supported by R01 GM123260 (M.R.P.), the Ruth L. Kirschstein National Research Service Award Individual Predoctoral Fellowship 1F31GM125344 (B.L.G.), and the Vanderbilt University Medical Center Diabetes Research and Training Center Pilot and Feasibility Grant. Confocal microscopy imaging was performed through the use of the Vanderbilt Cell Imaging Shared Resource (supported by NIH grants CA68485, DK20593, DK58404, DK59637 and EY08126). Quantification of mtDNA copy number and ΔmtDNA frequency was conducted with the help of the Simon A. Mallal Laboratory at Vanderbilt University Medical Center.

## Author contributions

Conceptualization, B.L.G. and M.R.P; Methodology, B.L.G, A.T.T., and M.R.P.; Formal Analysis, B.L.G, A.T.T., and M.R.P.; Investigation, B.L.G.; Resources, M.R.P.; Writing – Original Draft, B.L.G.; Writing – Review & Editing, B.L.G, A.T.T., and M.R.P.; Visualization, B.L.G.; Supervision, M.R.P.; Funding Acquisition, B.L.G. and M.R.P.

## STAR★ METHODS

### KEY RESOURCES TABLE

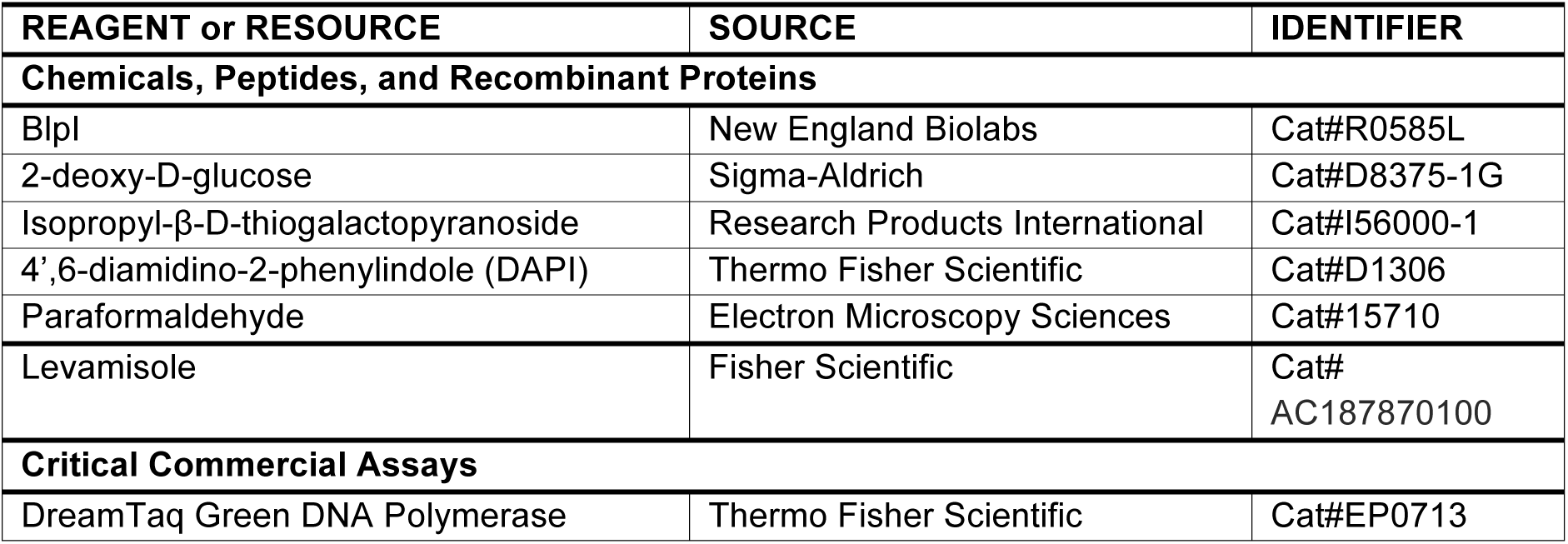
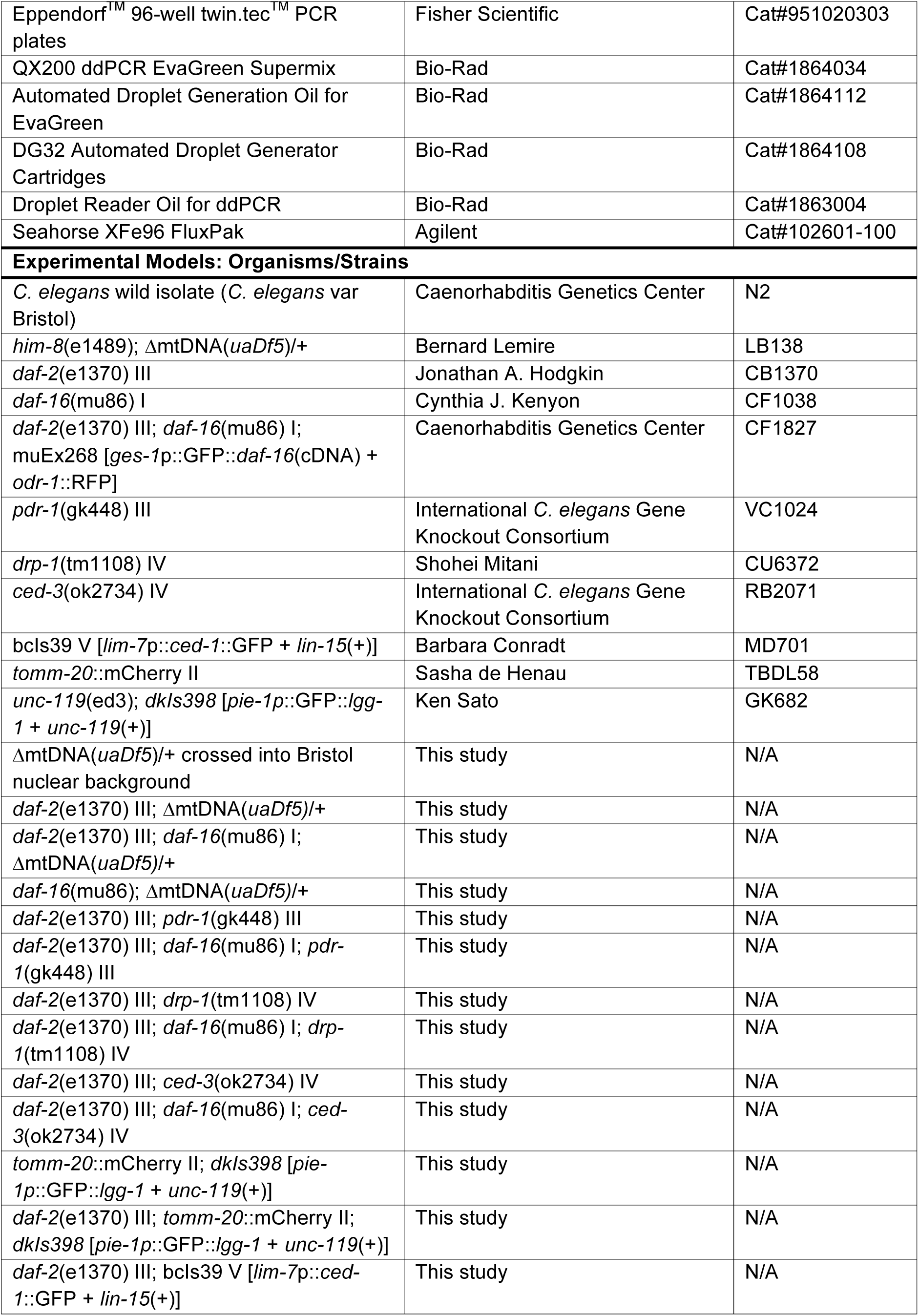
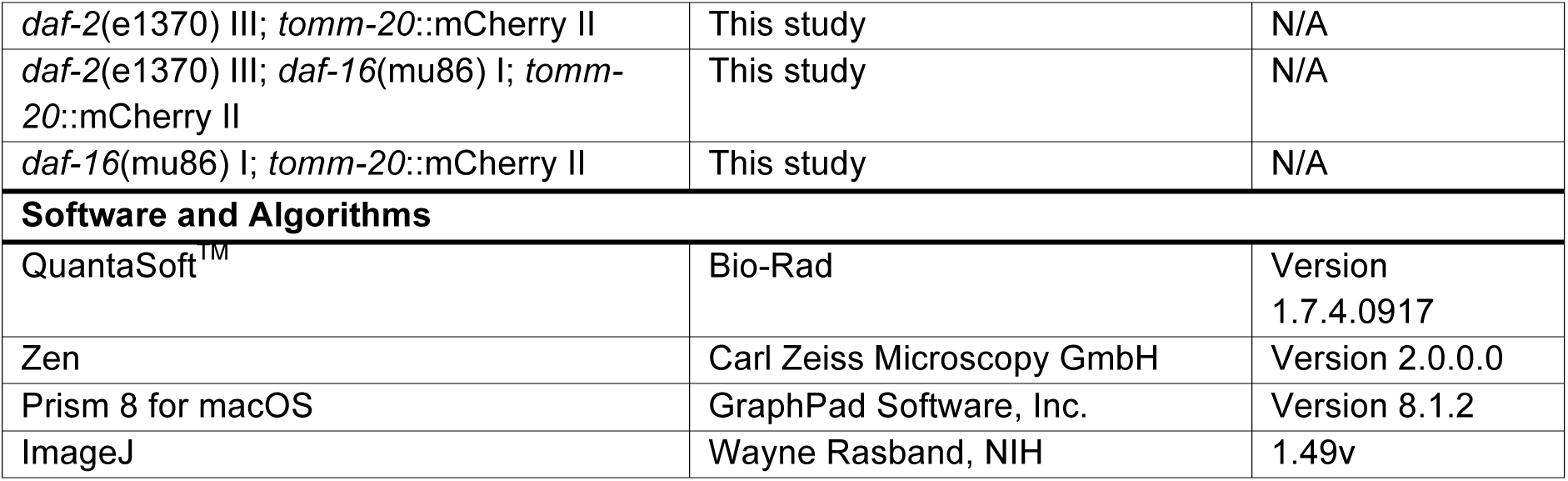

### EXPERIMENTAL#MODEL AND SUBJECT DETAILS

#### Nematode culture

*C. elegans* strains used in this study were maintained on 60-mm standard nematode growth medium (NGM) plates seeded with live OP50 *E. coli* bacteria as a food source, unless otherwise indicated in the METHOD DETAILS below. Nematode strains were incubated at 20°C unless otherwise indicated. Age-matched nematodes were used in all experiments with the exception of the multigenerational competition experiment (see below).

### METHOD DETAILS

#### Nematode lysis

To prepare nematodes for genotyping and quantification of mtDNA copy number and ΔmtDNA frequency, nematodes were lysed using the following protocol. Nematodes were transferred to sterile PCR tubes or 96-well PCR plates containing lysis buffer with 100 µg/mL proteinase K. Volume of lysis buffer varied by worm count: 10 µL for individual adults, pooled larvae, or pooled embryos; 20 µL for 5 or 10 pooled adults; 50 µL for pooled nematodes of mixed age (competition experiments, see below). Each tube or plate was then incubated at −80°C for 10 minutes, then at 60°C for 60 minutes (90 minutes for pooled nematodes), and then at 95°C for 15 minutes to inactivate the proteinase K. Nematode lysates were then stored at −20°C.

#### Genetic crosses and genotyping

To control for nuclear effects on ΔmtDNA proliferation, hermaphroditic nematodes carrying the ΔmtDNA allele *uaDf5* were serially back-crossed into a male stock of the Bristol (N2) *C. elegans* nuclear background for six generations. To investigate the role of insulin signaling in selfish mitochondrial genome dynamics, the alleles *daf-2*(e1370) and *daf-16*(mu86) were introduced to the ΔmtDNA heteroplasmic lineage by classical genetic crosses. To investigate the mechanistic basis by which the insulin signaling pathway regulates mtDNA levels, mutant alleles affecting various putative downstream processes were genetically crossed into the insulin signaling-defective nuclear genotypes. Specifically, the parkin-dependent mitophagy-defective *pdr-1*(gk448), the mitochondrial fission-defective *drp-1*(tm1108), and the apoptosis-defective *ced-3*(ok2734) were each genetically combined with *daf-2*(e1370), both with and without the *daf-16*(mu86) allele. Nuclear genotype was confirmed by PCR using the following oligonucleotide primers:

Mutant and wildtype mtDNA:

Exterior forward: 5’-CCATCCGTGCTAGAAGACAA-3’

Interior forward: 5’-TTGGTGTTACAGGGGCAACA-3’

Reverse: 5’-CTTCTACAGTGCATTGACCTAGTC-3’

*daf-2*:

Forward: 5’-CATCAAGATCCAGTGCTTCTGAATCGTC-3’

Reverse: 5’-CGGGATGAGACTGTCAAGATTGGAG-3’

*daf-16*:

Forward: 5’-CACCACGACGCAACACACTAATAGTG-3’

Exterior reverse: 5’-CACGAGACGACGATCCAGGAATCG-3’

Interior reverse: 5’-GGTCTAAACGGAGCAAGTGGTTACTG-3’

*pdr-1*:

Exterior forward: 5’-GAATCATGTTGAAAATGTGACGCGAG-3’

Interior forward: 5’-CTGACACCTGCAACGTAGGTCAAG-3’

Reverse: 5’-GATTTGACTAGAACAGAGGTTGACGAG-3’

*drp-1*:

Forward: 5’-CGTCGGATCACAGTCGGC-3’

Reverse: 5’-GCACTGACCGCTCTTTCTCC-3’

*ced-3*:

Exterior forward: 5’-CAGTACTCCTTAAAGGCGCACACC-3’

Interior forward: 5’-GATTGGTCGCAGTTTTCAGTTTAGAGGG-3’

Reverse: 5’-CGATCCCTGTGATGTCTGAAATCCAC-3’

The insulin signaling receptor allele *daf-2*(e1370) introduces a point mutation that eliminates a *BlpI* restriction endonuclease recognition site. Following PCR amplification, *daf-2* PCR products were incubated with *BlpI* and New England BioLabs CutSmart® buffer at 37°C for 2 hours prior to gel electrophoresis. Fluorescent reporters used in this study were genotyped by fluorescence microscopy.

#### Quantification of mtDNA copy number and ΔmtDNA frequency

Quantification of mtDNA copy number and ΔmtDNA frequency was accomplished using droplet digital PCR (ddPCR). Nematodes were lysed as described above. Lysates were then diluted in nuclease-free water, with a dilution factor varying depending on nematode concentration: 20x for embryos, 200x for pooled larvae, 200x for single adults, 1000x for pooled adults, 20,000x for pooled nematodes of mixed age from the competition experiments (control diet) or 2,000x for pooled nematodes of mixed age from the competition experiments (restricted diet). The lower dilution factor for the lysates collected from the restricted diet condition was due to the smaller population sizes of nematodes raised on a restricted diet, which arises from reduced fecundity under diet restriction and was reflected in the number of nematodes present in these lysates. Next, either 2 µL or 5 µL of each dilute nematode lysate was combined with 0.25 µL of a 10-µM aliquot of each of the following oligonucleotide primers:

For quantifying wildtype mtDNA:

5’-GTCCTTGTGGAATGGTTGAATTTAC-3’

5’-GTACTTAATCACGCTACAGCAGC-3’

For quantifying ΔmtDNA:

5-‘CCATCCGTGCTAGAAGACAAAG-3’

5-‘CTACAGTGCATTGACCTAGTCATC-3’

Mixtures of dilute nematode lysate and primer were combined with nuclease-free water and Bio-Rad QX200^TM^ ddPCR^TM^ EvaGreen Supermix to a volume of 25 µL in Eppendorf^TM^ 96-well twin.tec^TM^ PCR plates. Droplet generation and PCR amplification were performed according to manufacturer protocol with an annealing temperature of 58°C. For amplification of heteroplasmic nematode lysates, wildtype and ΔmtDNA primers were combined in the same reaction, and each droplet was scored as containing either wildtype or mutant template using the 2D amplitude (dual-wavelength) clustering plot option in the Bio-Rad QuantaSoft^TM^ program.

#### Respiration assay

Basal and maximum oxygen consumption rates were measured using the Seahorse XFe96 Analyzer in the High Throughput Screening Facility at Vanderbilt University. One day before experimentation, each well of a 96-well sensor cartridge that comes as part of the Seahorse XFe96 FluxPak was incubated with 200 µL of the Seahorse XF Calibrant Solution. On the day of the experiment, 10-20 L4-stage animals were placed into each well of the cell culture microplate. Wells contained either M9 buffer or 10 µM FCCP. After calibration, 16 measurements were performed at room temperature. Measurements 12 through 16 were averaged and normalized to number of worms per well.

#### Fertility

To assay fertility, day-2 adult nematodes were individually transferred onto NGM plates seeded with live OP50 *E. coli* and incubated at 20°C for 4 hours. The adults were then individually lysed as described above. Fertility was scored as the average number of viable progeny produced per hour during the 4-hour window, where viable progeny were identified as those that had progressed from embryos to larvae within 24 hours of being laid. The ΔmtDNA frequency of each parent was determined using ddPCR as described above.

#### Development

The impact of ΔmtDNA levels on development was assayed by comparing ΔmtDNA frequency with developmental stage for each nematode in a population of age-synchronized larvae. To age-synchronize larvae, multiple mature heteroplasmic adults carrying ΔmtDNA in the Bristol nuclear background were transferred to an NGM plate seeded with live OP50 *E. coli* and allowed to lay eggs at 20°C for 2 hours. Adults were then removed from the plate. After 48 hours, each nematode was individually lysed and its respective larval stage (L2, L3, or L4) was annotated. None of the nematodes had yet reached adulthood at this point. Embryos that failed to transition to larvae were discarded. The ΔmtDNA frequency of each larval nematode was determined using ddPCR as described above.

#### Sub-organismal selection assay

Sub-organismal selection for ΔmtDNA was quantified by measuring changes in ΔmtDNA frequency as a function of developmental stage, and as a function of initial (parental) ΔmtDNA frequency, within a single generation. This was accomplished using two complementary approaches. In the first approach, three individual age-synchronized parents were selected according to initial ΔmtDNA frequency (parents with low, middle, and high frequency). One age-matched (L4-stage) nematode was selected at random from each line respectively maintained under artificial selection for low (<50%), medium (50-70%), and high (>70%) ΔmtDNA frequency. Each of these nematodes was placed onto a fresh NGM plate seeded with live OP *E. coli* and incubated for 2 days at 20°C. Each day-2 adult was then transferred to a fresh food plate every 4 hours and allowed to lay embryos. At each 4-hour time point, approximately one third of the embryos produced were individually lysed. After 12 hours, the adults were individually lysed. A 12-hour time window for embryo production was chosen in order to generate a sufficient offspring count to allow for the establishment of single-brood frequency distributions of ΔmtDNA. The 12-hour time window was divided into 4-hour segments in order to maintain age-synchronicity, as each larva was lysed within 4 hours of being laid across the entire 12-hour period. After 2 days at 20°C, approximately one third of the L4-stage larvae were individually lysed in the same 4-hour segments to maintain age synchronicity. After an additional 2 days at 20°C, the remaining one third of offspring were individually lysed in 4-hour segments, as they reached the same age at which their respective parent was lysed. The ΔmtDNA frequency of each individual was determined using ddPCR as described above and a ΔmtDNA frequency distribution was generated for each offspring life stage.

In the second approach, multiple L4-stage heteroplasmic nematodes were selected at random from the stock of nematodes carrying ΔmtDNA in the Bristol nuclear background. These larvae were transferred to a fresh food plate and incubated for 2 days at 20°C. The day-2 adults were then segregated onto individual plates and incubated for 4 hours at 20°C to produce age-synchronized progeny. After 4 hours, each parent was individually lysed. Three embryos from each parent were also lysed at the same time, in one pooled lysate per three same-parent embryos. After 2 days, three L4-stage larvae were pooled and lysed from each parent, similar to the lysis of embryos. After another 2 days, three adult progeny were pooled and lysed from each parent as they reached the age at which the parents were lysed. Each parent-progeny lineages was individually segregated from the rest. Since ΔmtDNA impacts fecundity, the progeny from parents on the lower end of the ΔmtDNA frequency are expected to be overrepresented in the offspring sampled from a mixed cohort of parents. Lineages were therefore segregated to ensure that the ΔmtDNA frequency from each progeny lysate was being compared with that of its own respective parent, in order to minimize the effect of organism-level selection on ΔmtDNA. In addition, progeny from each time-point were lysed in pools of three to reduce the effect of random drift on ΔmtDNA frequency. The ΔmtDNA frequency of parents and each developmental stage of progeny was determined using ddPCR as described above. For the measurement of sub-organismal selection on ΔmtDNA under nutrient-variable conditions, each parent was raised from embryo to adult under its respective dietary condition (diet restriction or control).

### Experimental evolution (organismal selection)

Selection against ΔmtDNA that occurs strictly at the level of organismal fitness was measured using a competition assay. Heteroplasmic nematodes carrying ΔmtDNA in the Bristol nuclear background were combined with Bristol-strain nematodes on 10-cm NGM plates seeded with live OP50 *E. coli*. For the first generation, heteroplasmic and Bristol strain nematodes were age-synchronized. Age synchronization was accomplished using a bleaching protocol. Nematodes from a mixed-age stock food plate were washed off the plate and into a sterile 1.7 mL microcentrifuge tube with nuclease-free water. The water was brought to a volume of 750 µL. The volume of each tube was brought to 1 mL by adding 100 µL of 5 N NaOH and 150 µL of 6% bleach. Each nematode tube was incubated at room temperature for 10 minutes with light vortexing every 2 minutes to rupture gravid adults and release embryos. Nematode tubes were centrifuged for 1 minute at 1,000x g to pellet the nematode embryos. To wash the nematode pellets, the supernatant was removed and replaced with 1 mL of nuclease-free water. After a second spin for 1 minute at 1,000x g, the water was removed and the nematode embryos were resuspended in 100 µL M9 buffer. The resuspended embryos were then transferred to glass test tubes containing 500 µL M9 buffer and incubated overnight at room temperature on a gentle shaker to allow hatching and developmental arrest at the L1 larval stage. On the following day, a glass Pasteur pipette was used to transfer approximately equal quantities of heteroplasmic and homoplasmic-wildtype nematodes onto the 10-cm food plates. Approximately 500 nematodes were transferred to each plate. In addition to 8 competition lines, 8 control lines were established by transferring only heteroplasmic nematodes onto food plates, with no homoplasmic-wildtype nematodes to compete against.

Every 3 days, the generation for each experimental line was reset. To do this, nematodes were washed off the plates using sterile M9 buffer into a sterile 1.7 mL collection tube. Approximately 500 nematodes of mixed age from each line were transferred to a fresh food plate. An additional 500 nematodes were lysed together in a single pooled lysate. Finally, 48 additional adults from each competition line were lysed individually in order to determine the fraction of heteroplasmic nematodes in each competition line at each generational time point. This experiment was continued for 10 consecutive generations.

Experimental evolution was also carried out to quantify nutrient-conditional organism-level selection. These conditions included 10-cm NGM plates seeded with a restricted or a control diet consisting of UV-killed OP50 *E. coli* (prepared as described below). Two iterations of this experiment were conducted, one with wildtype nuclear genotype and one with nematodes homozygous for the null *daf-16*(mu86) allele. Due to the smaller brood sizes among nematodes raised on a restricted diet, 200 nematodes were transferred and another 200 lysed at each generation, instead of the 500 as in the case of the experiment using a live bacterial diet. For these nutrient-conditional competition experiments, 6 replicate lines were propagated for each condition for a total of 8 consecutive generations. Lysis and quantification of ΔmtDNA frequency by ddPCR were performed as described above.

### Diet restriction

Diet restriction was accomplished using variable dilutions of UV-inactivated OP50 *E. coli* bacterial lawns on NGM plates. To prepare diet-restricted food plates, 1 L of sterile 2xYT liquid microbial growth medium was inoculated with 1 mL of live OP50 *E. coli* (suspended in liquid LB) using a sterile serological pipette. The inoculated culture was then incubated overnight on a shaker at 37°C. The following day, the OP50 *E. coli* was pelleted by centrifugation for 6 minutes at 3,900 rpm. The pellet was resuspended to a bacterial concentration of approximately 2×10^10^ cells/mL in sterile M9 buffer. This suspension was seeded onto NGM plates (control) or further diluted 100-fold to 2×10^8^ cells/mL in sterile M9 buffer before being seeded onto NGM plates (diet restriction). Plates were incubated upright at room temperature 4 hours to allow the lawns to dry. To inhibit bacterial growth, plates were irradiated with UV radiation using a Stratagene® UV Stratalinker 1800 set to 9.999×10^5^ µJ/cm^2^. To confirm inhibition of bacterial growth, UV-treated plates were incubated overnight at 37°C.

### Insulin signaling inactivation

Insulin signaling was conditionally inactivated using the allele *daf-2*(e1370), encoding a temperature-sensitive variant of the *C. elegans* insulin receptor homolog. Because complete loss of insulin signaling during early larval development results in a stage of developmental arrest (dauer), age-synchronized nematodes were incubated at the permissive temperature of 16°C until reaching the fourth and final larval stage. L4-stage larvae were then selected at random for either transfer to the restrictive temperature of 25°C or for continued incubation at 16°C as a control. After 4 days of incubation, mature adults were lysed and ddPCR quantification of ΔmtDNA frequency was performed as described above. To follow up on the downstream mechanism by which insulin signaling regulates mtDNA dynamics, homoplasmic nematodes were incubated at the restrictive temperature of 25°C and mtDNA copy number was measured using the same ddPCR primer pair that was used for quantifying the wildtype mtDNA in heteroplasmic worms.

### Knockdown of gene expression

Expression knockdown of the *C. elegans* insulin signaling receptor homolog, *daf-2*, was accomplished using feeder plates. Cultures consisting of 2 mL LB and 10 µL ampicillin were inoculated with a bacterial culture obtained from Source BioScience harboring the Y55D5A_391.b (*daf-2*) ORF plasmid clone and incubated overnight on a shaker at 37°C. Bacteria containing the empty plasmid vector were used to establish a control diet. The following day, 750 µL of culture was transferred to a flask containing 75 mL LB and 375 µL ampicillin and incubated 4-6 hours on a shaker at 37°C, until OD_550-600_ >0.8. An additional 75 mL LB was added to the culture along with another 375 µL ampicillin and 600 µL 1 M isopropyl β-D-1-thiogalactopyranoside (IPTG) to induce expression of the small interfering RNA. Cultures were incubated another 4 hours on a shaker at 37°C. Cultures were then centrifuged for 6 minutes at 3,900 rpm and the resulting bacterial pellets were each resuspended in 6 mL M9 buffer with 8 mM IPTG. Next, 250 µL of resuspension was seeded onto each NGM plate. Plates were allowed to dry at room temperature in the dark and then stored at 4°C until use. Synchronized L4-stage nematodes were transferred at random to either RNAi knockdown or control plates and incubated at 25°C until day 4 of adulthood to match the conditions that were used for the *daf-2* mutant allele. Day-4 adults were lysed and their mtDNA copy number was quantified using ddPCR as described above.

### Live imaging

Overall mitochondrial content across the wildtype and defective insulin signaling genotypes was measured using the mitochondrial reporter TOMM-20::mCherry. Age-synchronized nematodes were incubated for 2 days from the L4 stage to mature adulthood at 25°C, immobilized with 10 mM levamisole, and placed on the center of a 2% agarose pad on a microscope slide. Nematodes were imaged at 10x magnification using a Leica DM6000 B compound fluorescence microscope and mitochondrial fluorescence was quantified using ImageJ. Apoptosis was imaged in *daf-2*(e1370) mutant nematodes and wildtype controls using the CED-1::GFP reporter. Age-synchronized nematodes were incubated for 2 days from the L4 stage to mature adulthood at 25°C before being immobilized and mounted on microscope slides as described above. Apoptotic cells were imaged using a Zeiss LSM 880 Confocal Laser Scanning microscope at 20x magnification.

### Staining and imaging of germline nuclei

Nematode germline nuclei were quantified across age-synchronized mature adults homozygous for *daf-2*(e1370) or *daf-16*(mu86), as well as in double-mutants and wildtype controls. For each genotype, age-synchronized L4-stage nematodes were incubated for 2 days at 25°C and then placed in a plate containing 3 mL of PBS with 200 µM levamisole. To dissect the nematode gonads, each adult was decapitated using two 25Gx1” hypodermic needles in a scissor-motion under a dissecting microscope. Dissected gonads were fixed for 20 minutes in 3% paraformaldehyde. Fixed gonads were transferred to a glass test tube using a glass Pasteur pipette and the paraformaldehyde was replaced with PBT (PBS buffer with 0.1% Tween 20) and incubated for 15 minutes at room temperature. The PBT was then replaced with PBT containing 100 ng/mL 4’,6’-diamidino-2-phenylindole dihydrochloride (DAPI) and the gonads were incubated in darkness for another 15 minutes at room temperature. Gonads were then subjected to 3x consecutive washes, each consisting of a 1-minute centrifugation at 1,000 rpm followed by replacement of the PBT. Gonads were then mounted directly onto a 2% agarose pad on the center of a microscope slide and imaged using a Zeiss LSM 880 Confocal Laser Scanning microscope at 20x magnification.

---

**Figure S1, related to Figure 1.**
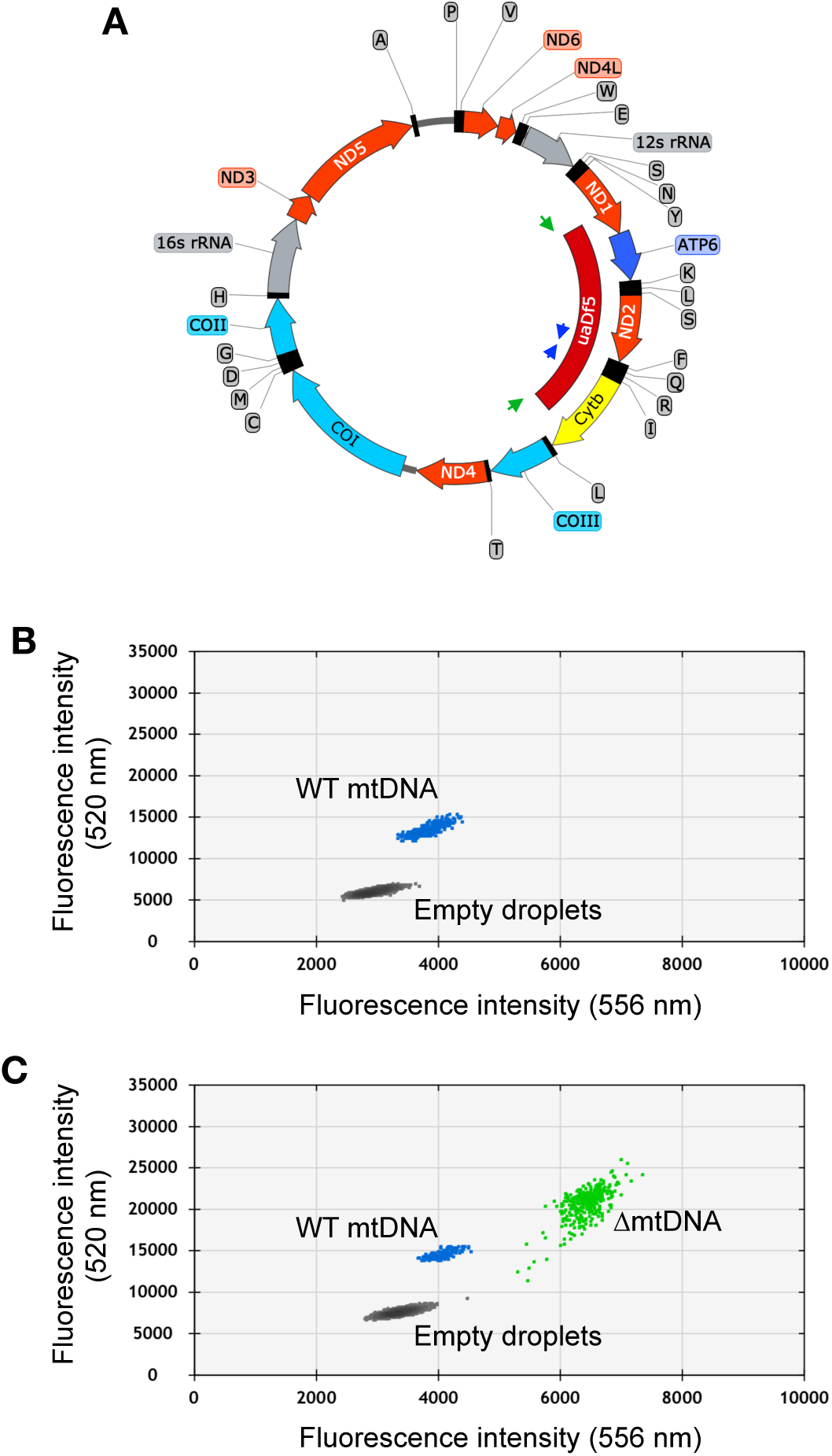
Quantification of mtDNA copy number and ΔmtDNA frequency by droplet digital PCR. (A) Map of mtDNA showing the *uaDf5* deletion (ΔmtDNA) and the strategy for oligonucleotide primer design for the multiplex quantification of ΔmtDNA and wildtype mtDNA simultaneously within the same reaction. Due to the deletion size, primers flanking the *uaDf5* deletion (green arrows) amplify a PCR product off of the ΔmtDNA but not wildtype mtDNA template (see panel B). Likewise, primers complementary to a sequence within the region spanning the *uaDf5* deletion (blue arrows) amplify a PCR product off of the wildtype mtDNA but not ΔmtDNA template. (B-C) Sample droplet digital PCR data plots showing mtDNA copy number in lysates from homoplasmic wildtype (B) and heteroplasmic (C) nematodes. Mutant frequency is determined from ΔmtDNA copy number over total copy number.

**Figure S2, related to Figure 2.**
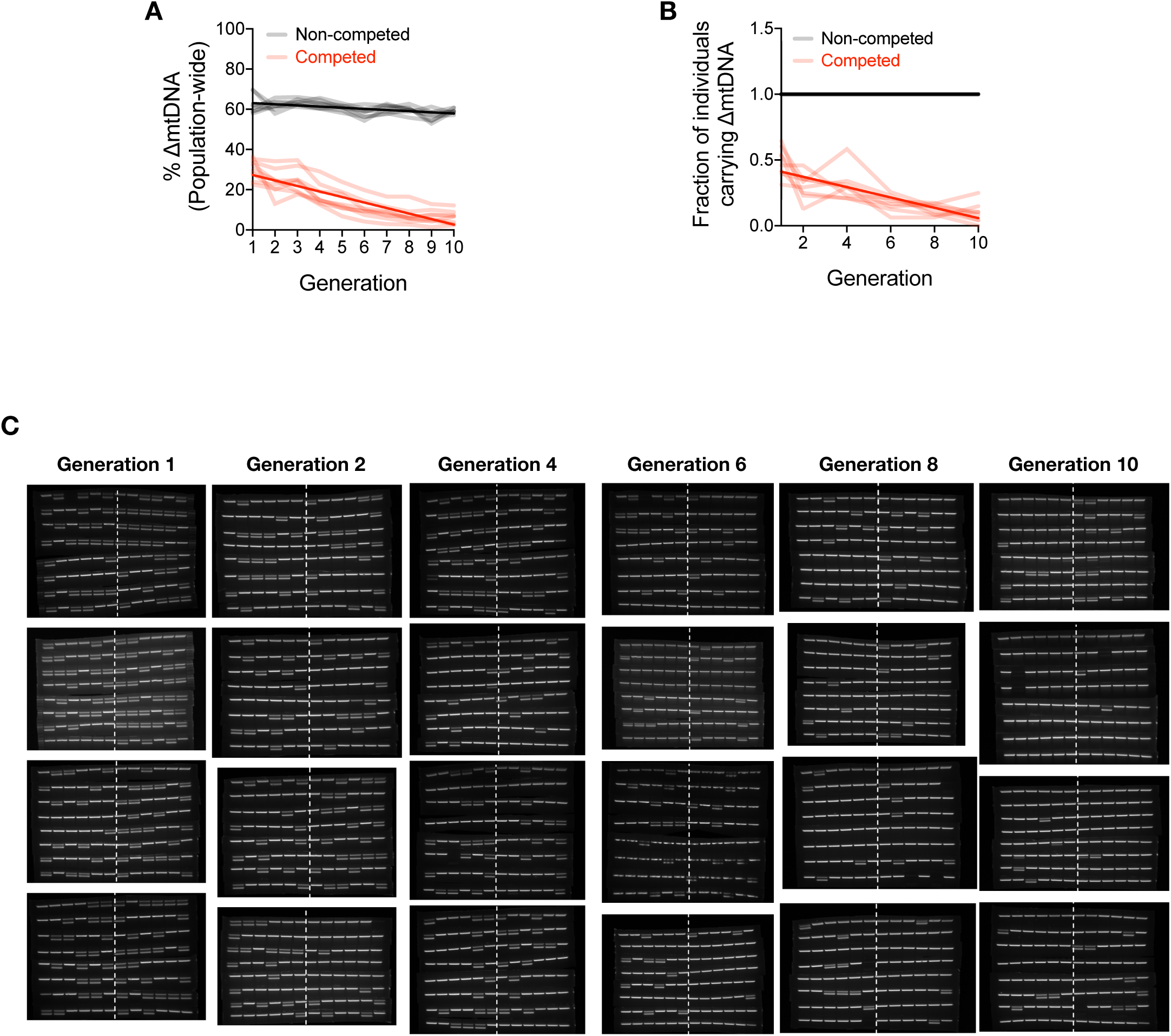
Change in ΔmtDNA frequency in competition experiments is due to organismal selection. (A) Quantification of ΔmtDNA frequency across 8 replicate competed (red) and non-competed (black) lineages. Same experiment as shown in Figure 2A but with y-axis expressing raw ΔmtDNA frequency measurements. Dark lines reflect best-fit regression across all replicate lineages. (B) Quantification of fraction of heteroplasmic members from the competition experiment shown in (A), based on gel images shown (C). (C) PCR gel images reflecting individual adult nematodes sampled from the competition lines in the organism-level selection experiment. Approximately 48 adults were individually lysed from the first generation and every second generation thereafter. A single PCR band reflects Bristol strain nematodes (homoplasmic for wildtype mtDNA), whereas double bands reflect heteroplasmic animals carrying the ΔmtDNA allele. The numbers to the sides of the gel images label the replicate lineages.

**Figure S3, related to Figure 3.**
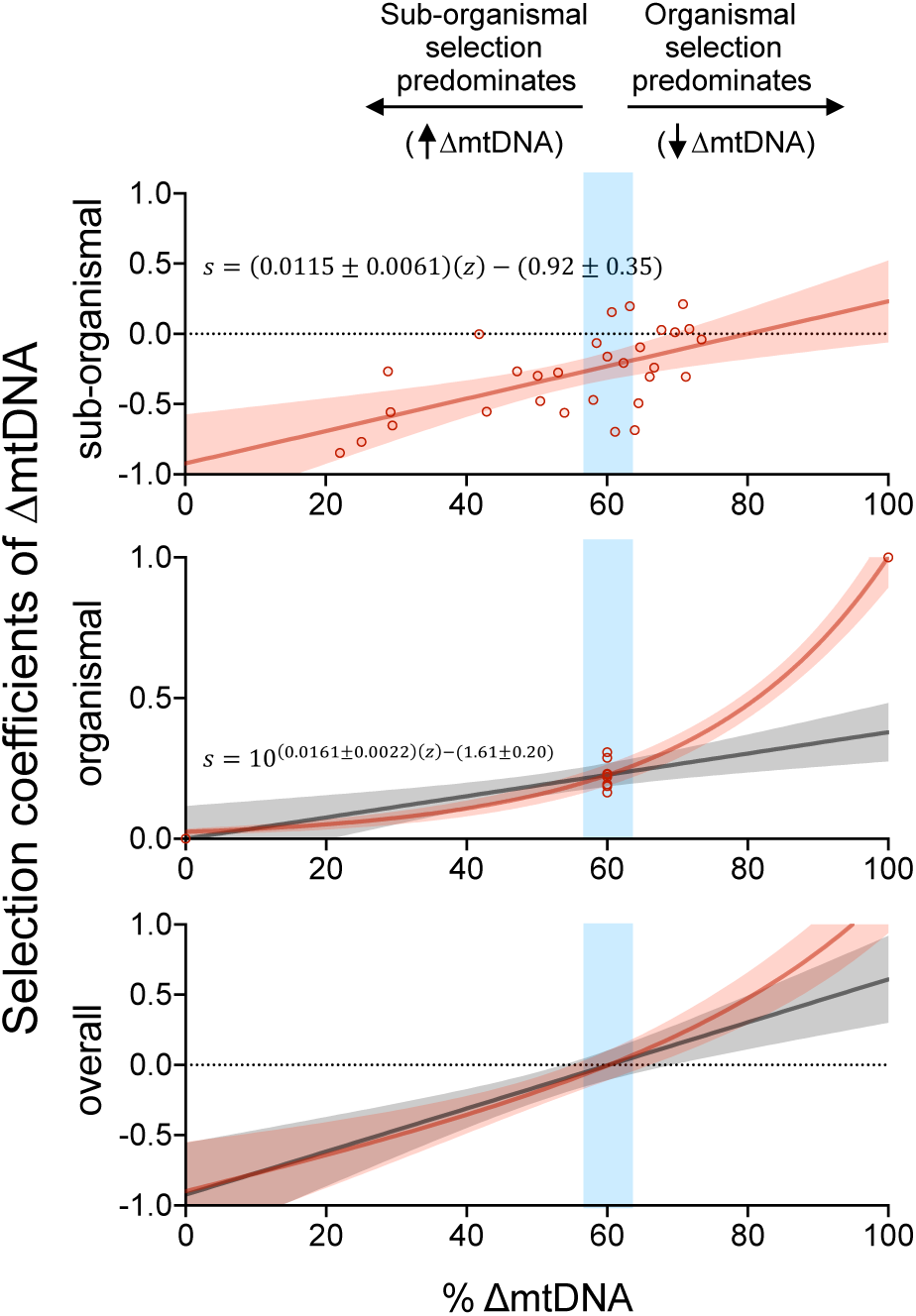
Fitness of wildtype mtDNA relative to ΔmtDNA as a function of ΔmtDNA frequency, at each selection level, assuming organismal selection increases linearly with ΔmtDNA frequency. Selection coefficient against ΔmtDNA at sub-organismal (top), organismal (middle), and overall combined (bottom) levels of selection, similar to Figure 3 but assuming that the strength of organismal selection increases linearly with ΔmtDNA frequency. The linear regression (gray) and the non-linear regression (red, also shown in Figure 3) of frequency-dependent organismal selection against ΔmtDNA predict that sub-organismal and organismal selection cancel out near 60%, allowing ΔmtDNA to persist in this range.

**Figure S4, related to Figure 3.**
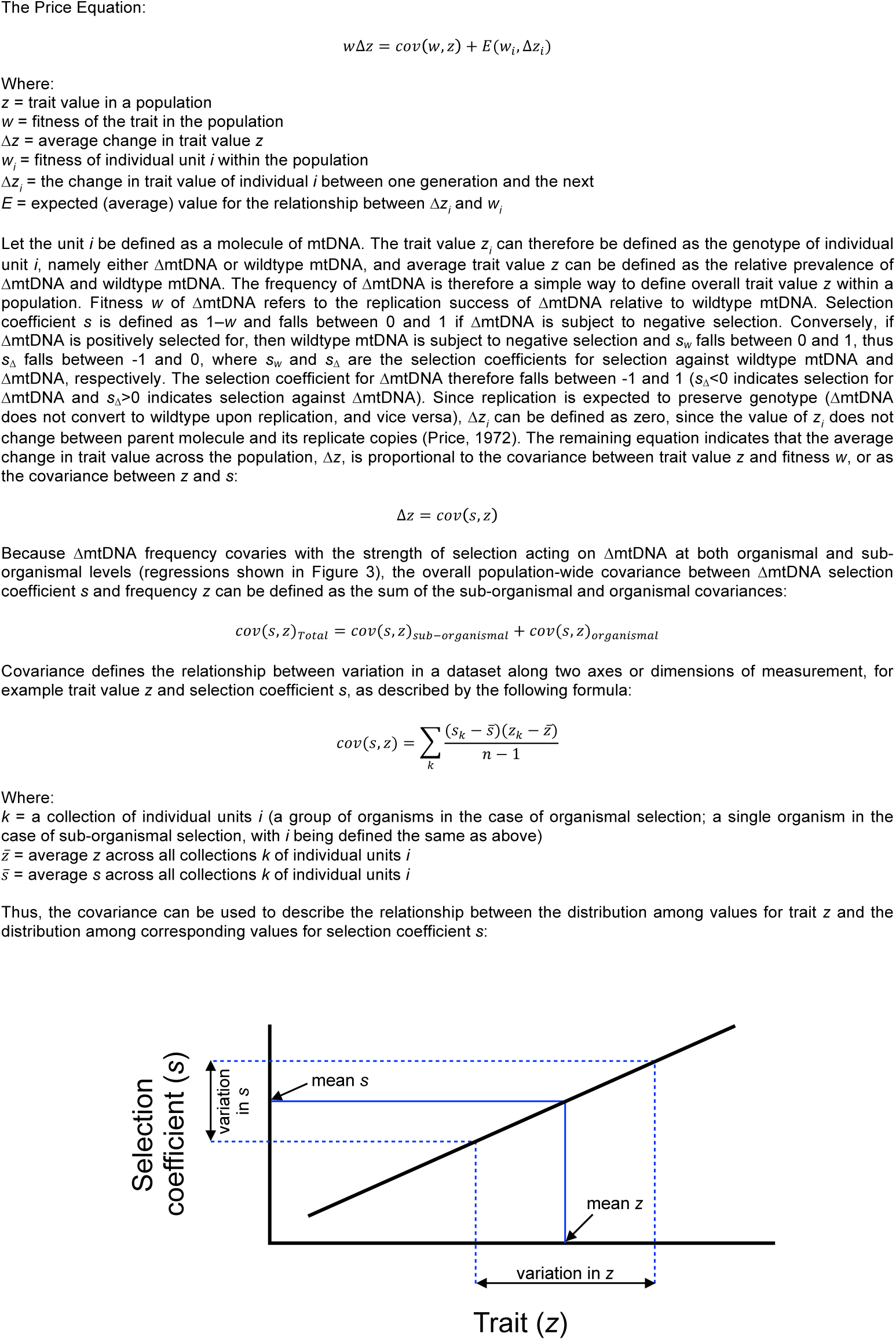
Description of the Price Equation and trait-fitness covariance. The Price Equation (top) shows that the change in a trait value (Δ*z*) is proportional to the covariance between the trait value itself and its fitness. Covariance describes the relationship between the variation in one value (e.g. trait) and variation in a separate value (e.g. fitness), assuming the two sets of values are correlated (bottom graph).

**Figure S5, related to Figure 4.**
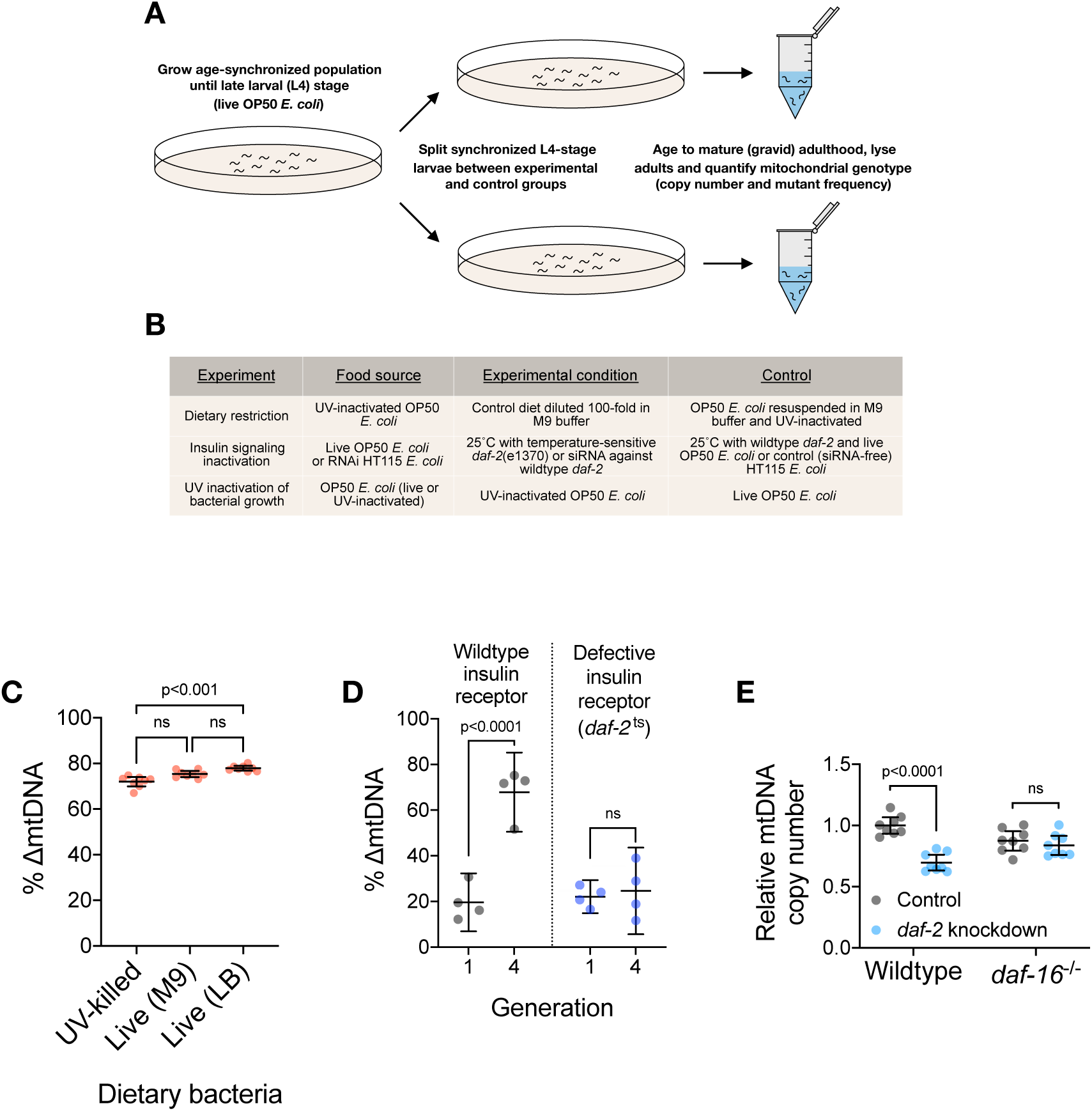
Diet and insulin signaling regulate mtDNA copy number and ΔmtDNA frequency. (A) Schematic of experimental workflow for assaying the impact of dietary and pharmacological perturbations on ΔmtDNA frequency. (B) Experimental conditions under which changes in ΔmtDNA frequency were assayed. (C) The frequency of ΔmtDNA in age-synchronized, pooled adults (10 adults per sample) grown on one of three food plates: UV-killed OP50 *E. coli*, live OP50 suspended in M9 buffer, and live OP50 suspended in LB medium. Kruskal-Wallis ANOVA with Dunn’s multiple comparisons test. (D) ΔmtDNA frequency in generation 1 versus generation 4, between wildtype and the temperature-sensitive allele *daf-2*(e1370). Two-way ANOVA with Bonferroni correction. N=4 lysates containing 5 pooled age-synchronized adults each. (E) mtDNA copy number in age-synchronized adults lacking ΔmtDNA and expressing either wildtype or null *daf-16*(mu86), raised on either a *daf-2* RNAi knockdown diet or an empty-vector control diet. Two-way ANOVA with Bonferroni correction. N=8 lysates containing 5 pooled age-synchronized. Error bars represent 95% C.I.

**Figure S6, related to Figure 5.**
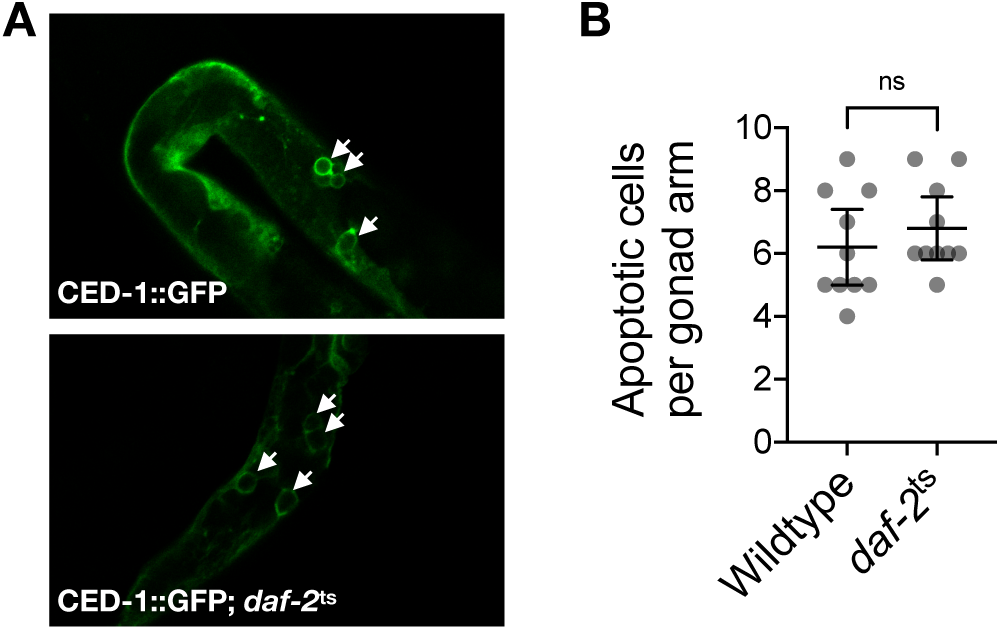
No observed change in germline apoptosis between wildtype and *daf-2* mutants. Images (A) and quantification (B) of apoptosis as indicated by the engulfment of apoptotic cells by the reporter CED-1::GFP (white arrows in A), between age-synchronized adults of wildtype or temperature-sensitive *daf-2*(e1370) mutant genotype. Mann-Whitney test. Error bars represent 95% C.I.

**Figure S7, related to Figure 7.**
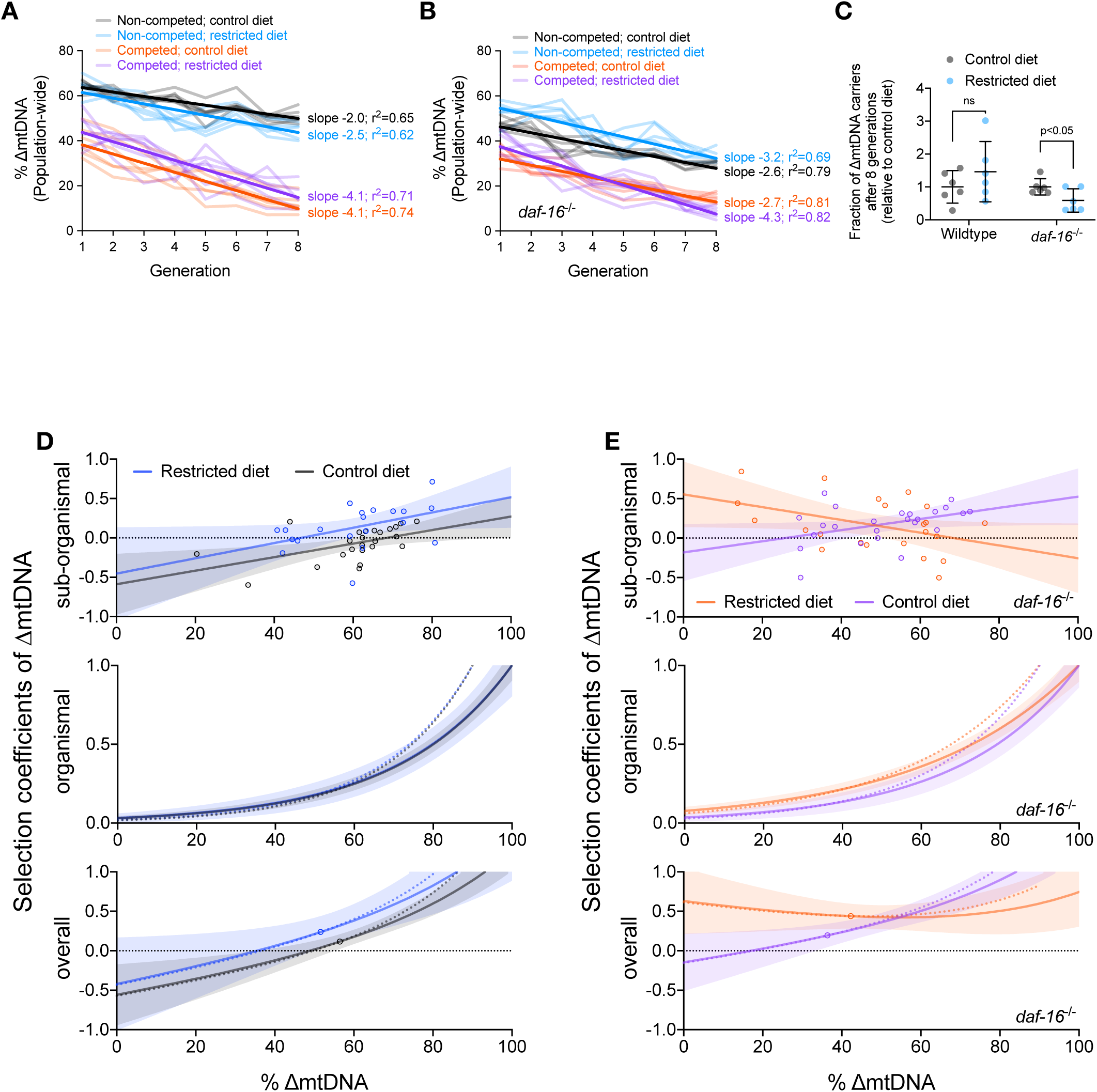
Quantification of multilevel selection between dietary conditions and *daf-16* genotypes. (A-B) Same data presented in Figure 7C-7E but with y-axis representing raw ΔmtDNA frequency measurements. Dark lines reflect best-fit regression across all replicate lineages. (C) Fraction of ΔmtDNA-carrying individuals at generation 8 of the competition experiment shown in (A) and (B), normalized to control-diet lines. (D-E) Selection coefficients against ΔmtDNA at sub-organismal (top graphs), organismal (middle graphs), and overall combined (bottom graphs) levels of selection, in restricted versus control diets, in animals with either a wildtype (D) or null *daf-16*(mu86) (E) nuclear genotype. Shaded regions represent 95% C.I. Dotted lines (middle and lower graphs) show regressions if selection coefficient is 1.0 at 90% ΔmtDNA frequency (confidence intervals omitted from these regressions for simplicity). Open-circle data points on overall regressions (bottom graphs) correspond to ΔmtDNA frequency of non-competing lineages averaged across all 8 generations.

**Table S1.** Raw shifts in ΔmtDNA at sub-organismal and organismal levels.

| Selection level | Sub-organismal | Organismal |
| --- | --- | --- |
| **Units** |  |  |
| X-axis | % ΔmtDNA (parents) | Time (generations) |
| Y-axis | Shift in % ΔmtDNA<br>(progeny - parent) | Population-wide % ΔmtDNA<br>(Mean individual % ΔmtDNA: 60) |
| **Best-fit values** |  |  |
| Slope | -0.1975 | -2.743 |
| Intercept | 15.92 | 30.07 |
| **Std. Error** |  |  |
| Slope | 0.0546 | 0.206 |
| Intercept | 3.11 | 1.278 |
| **Goodness of Fit** |  |  |
| R square | 0.3183 | 0.6945 |
| Sy.x | 4.465 | 5.292 |
| **Linear regression statistics** |  |  |
| N | 30 | 80 |
| F | 13.08 | 177.3 |
| P (non-zero slope) | 0.0012 | <0.0001 |

**Table S2.** Selection coefficient at sub-organismal and organismal levels.

| Selection level | Sub-organismal | Organismal |
| --- | --- | --- |
| **Units** |  |  |
| X-axis | % ΔmtDNA (parents) | Mean heteroplasmic % ΔmtDNA |
| Y-axis | Selection coefficient | Selection coefficient |
| **Best-fit values** |  |  |
| Slope | 0.0115 | 0.0161 |
| Intercept | -0.92 | -1.61 |
| **Std. Error** |  |  |
| Slope | 0.0030 | 0.0009 |
| Intercept | 0.17 | 0.08 |
| **Goodness of Fit** |  |  |
| R square | 0.35 | 0.97 |
| Sy.x | 0.24 | 0.05 |
| **Linear regression statistics** |  |  |
| N | 30 | 8 |
| F | 14.9 | N/A |
| P (non-zero slope) | <0.001 | N/A |

**Table S3.** Raw shifts in ΔmtDNA frequency by diet, daf-16 genotype, and parental ΔmtDNA level.

| Genotype<br>Diet | Wildtype |  | *daf-16<sup>-/-</sup>* |  |
| --- | --- | --- | --- | --- |
|  | Control (*ad libitum*) | Restricted | Control (*ad libitum*) | Restricted |
| **Units** |  |  |  |  |
| X-axis | % ΔmtDNA (parents) | % ΔmtDNA (parents) | % ΔmtDNA (parents) | % ΔmtDNA (parents) |
| Y-axis | Shift in % ΔmtDNA<br>(progeny - parent) | Shift in % ΔmtDNA<br>(progeny - parent) | Shift in % ΔmtDNA<br>(progeny - parent) | Shift in % ΔmtDNA<br>(progeny - parent) |
| **Best-fit values** |  |  |  |  |
| Slope | -0.157 | -0.294 | -0.189 | 0.0978 |
| Intercept | 10.66 | 13.34 | 4.301 | -9.319 |
| **Standard Error** |  |  |  |  |
| Slope | 0.064 | 0.125 | 0.086 | 0.107 |
| Intercept | 3.971 | 7.78 | 4.494 | 5.5 |
| **Goodness of Fit** |  |  |  |  |
| R square | 0.214 | 0.234 | 0.181 | 0.0402 |
| Sy.x | 3.889 | 7.063 | 5.772 | 8.734 |
| **Linear regression statistics** |  |  |  |  |
| N | 24 | 20 | 24 | 22 |
| F | 5.992 | 5.5 | 4.845 | 0.8378 |
| P (non-zero slope) | 0.0228 | 0.0307 | 0.0385 | 0.3709 |

**Table S4.**
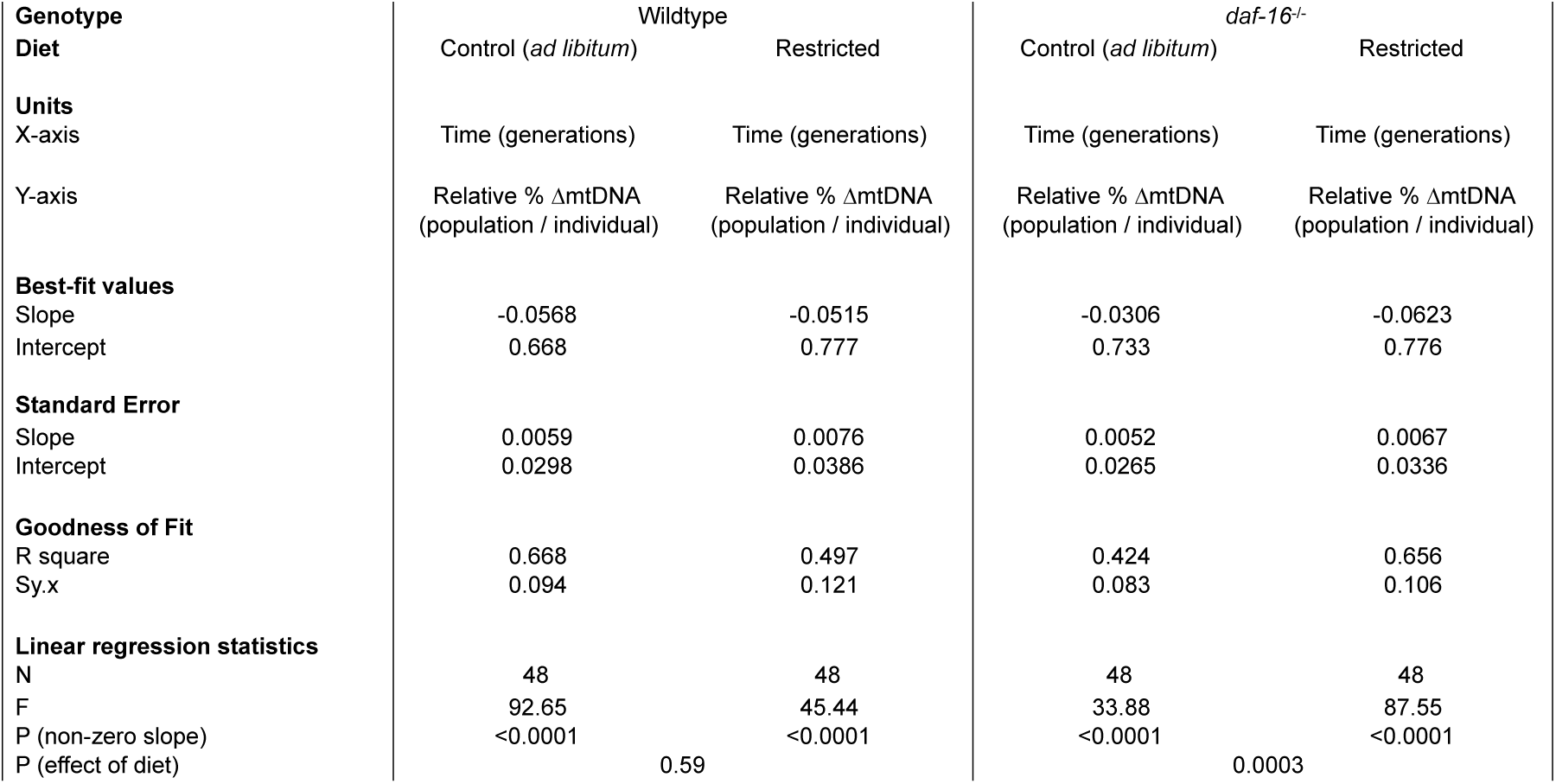
Organismal selection on ΔmtDNA by diet and *daf-16* genotype.

**Table S5.** Sub-organismal selection coefficients by diet, daf-16 genotype, and parental ΔmtDNA level.

| Genotype<br>Diet | Wildtype |  | *daf-16<sup>-/-</sup>* |  |
| --- | --- | --- | --- | --- |
|  | Control (*ad libitum*) | Restricted | Control (*ad libitum*) | Restricted |
| **Units** |  |  |  |  |
| X-axis | % ΔmtDNA (parents) | % ΔmtDNA (parents) | % ΔmtDNA (parents) | % ΔmtDNA (parents) |
| Y-axis | Selection coefficient | Selection coefficient | Selection coefficient | Selection coefficient |
| **Best-fit values** |  |  |  |  |
| Slope | 0.0086 | 0.0097 | 0.0071 | -0.0081 |
| Intercept | -0.59 | -0.45 | -0.18 | 0.55 |
| **Standard Error** |  |  |  |  |
| Slope | 0.0030 | 0.0045 | 0.0033 | 0.0039 |
| Intercept | 0.19 | 0.28 | 0.17 | 0.20 |
| **Goodness of Fit** |  |  |  |  |
| R square | 0.27 | 0.21 | 0.17 | 0.18 |
| Sy.x | 0.18 | 0.25 | 0.22 | 0.32 |
| **Linear regression statistics** |  |  |  |  |
| N | 24 | 20 | 24 | 20 |
| F | 8.16 | 4.66 | 4.48 | 4.36 |
| P (non-zero slope) | <0.01 | <0.05 | <0.05 | <0.05 |

**Table S6.** Organismal selection coefficients by diet, *daf-16* genotype, and ΔmtDNA level.

| Genotype<br>Diet | Wildtype |  | *daf-16<sup>-/-</sup>* |  |
| --- | --- | --- | --- | --- |
|  | Control (*ad libitum*) | Restricted diet | Control (*ad libitum*) | Restricted diet |
| **Units** |  |  |  |  |
| X-axis | Mean heteroplasmic % ΔmtDNA | Mean heteroplasmic % ΔmtDNA | Mean heteroplasmic % ΔmtDNA | Mean heteroplasmic % ΔmtDNA |
| Y-axis | Selection coefficient | Selection coefficient | Selection coefficient | Selection coefficient |
| **Best-fit values** |  |  |  |  |
| Slope | 0.0151 | 0.0149 | 0.0145 | 0.0112 |
| Intercept | -1.51 | -1.49 | -1.45 | -1.115 |
| **Standard Error** |  |  |  |  |
| Slope | 0.0008 | 0.0017 | 0.0014 | 0.0007 |
| Intercept | 0.08 | 0.16 | 0.13 | 0.06 |
| **Goodness of Fit** |  |  |  |  |
| R square | 0.98 | 0.94 | 0.97 | 0.98 |
| Sy.x | 0.04 | 0.08 | 0.06 | 0.04 |

